# Contribution of metal transporters of the ABC, ZIP, and NRAMP families to manganese uptake and infective endocarditis virulence in *Streptococcus sanguinis*

**DOI:** 10.1101/2021.05.28.446176

**Authors:** Tanya Puccio, Karina S. Kunka, Todd Kitten

## Abstract

*Streptococcus sanguinis* is an important cause of infective endocarditis. In strain SK36, the ABC-family manganese transporter, SsaACB, is essential for virulence. We have now identified a ZIP-family protein, TmpA, as a secondary manganese transporter. A *tmpA* mutant had no phenotype, but a Δ*ssaACB* Δ*tmpA* mutant was far more attenuated for serum growth and somewhat more attenuated for virulence in a rabbit model than its Δ*ssaACB* parent. The growth of both mutants was restored by supplemental manganese, but the Δ*ssaACB* Δ*tmpA* mutant required twenty-fold more and accumulated less. Although ZIP-family proteins are known for zinc and iron transport, TmpA-mediated transport of either metal was minimal. In contrast to *ssaACB* and *tmpA*, which appear ubiquitous in *S. sanguinis*, a *mntH* gene encoding an NRAMP-family transporter has been identified in relatively few strains, including VMC66. As in SK36, deletion of *ssaACB* greatly diminished VMC66 endocarditis virulence and serum growth, and deletion of *tmpA* from this mutant diminished virulence further. Virulence was not significantly altered by deletion of *mntH* from either VMC66 or its Δ*ssaACB* mutant. This and the accompanying paper together suggest that SsaACB is of primary importance for endocarditis virulence while secondary transporters TmpA and MntH contribute to growth under differing conditions.

## Introduction

Infective endocarditis (IE) is a disease caused by microorganisms entering the bloodstream and colonizing damaged heart valves, leading to potentially fatal complications such as congestive heart failure, aneurysm, and stroke (Bashore *et al.*, 2006). Many recent studies suggest that IE incidence is rising (Quan *et al.*, 2020), and morality rates of 12-30% are common (Jamil *et al.*, 2019, Ly *et al.*, 2020). Currently, prevention is limited to prophylactic antibiotics before dental procedures (Wilson *et al.*, 2021). The economic burden, potential for side effects, and questionable efficacy (Dayer & Thornhill, 2018, Thornhill *et al.*, 2018, Quan *et al.*, 2020) of this practice, as well as the increasing prevalence of antibiotic resistance (Dodds, 2017) all suggest the need for new approaches to prevention. In addition, the ability of oral bacteria to enter the bloodstream through any opening in the oral mucosa, which may occur during routine hygiene practices or mastication (Wray *et al.*, 2008, Wilson *et al.*, 2007), explains why prophylaxis given before dental procedures can never prevent all IE. Thus, it would be desirable to identify an inhibitor that is specific to endocarditis pathogens and which could be taken on a daily basis without disturbing the microbiome or selecting for resistance.

*Streptococcus sanguinis* is one of the most common oral bacteria to be isolated from IE patients (Di Filippo *et al.*, 2006). It is typically considered a commensal in the oral cavity due to an antagonistic relationship with the caries pathogen *Streptococcus mutans* (Kreth *et al.*, 2005). The trace element manganese (Mn) plays a role in the virulence of *S. sanguinis* (Das *et al.*, 2009, Crump *et al.*, 2014) as well as other streptococci (Eijkelkamp *et al.*, 2015) and bacterial pathogens (Juttukonda & Skaar, 2015, Kelliher & Kehl-Fie, 2016). Manganese import in bacteria has been confirmed in at least two protein families: ATP-binding cassettes (ABC) and natural resistance-associated macrophage proteins (NRAMP) (Waters, 2020). A knockout mutant of the lipoprotein component of an ABC manganese transporter in *S. sanguinis*, SsaB, is deficient in manganese transport as well as in aerobic serum growth (Crump *et al.*, 2014). This growth defect was rescued by the addition of only 2 μM Mn^2+^, indicating that manganese is able to enter *S. sanguinis* cells despite the absence of the primary transporter. The genome annotation of *S. sanguinis* SK36 (Xu *et al.*, 2007) did not indicate the presence of any canonical manganese transporters, which led us to examine other metal transport protein families. NRAMP-family proteins (Nevo & Nelson, 2006) are encoded by at least eight *S. sanguinis* strains but not by SK36. These proteins, often named MntH, have been found to contribute to manganese uptake and acid tolerance in other streptococci (Shabayek *et al.*, 2016, Kajfasz *et al.*, 2020).

The family of <u>Z</u>RT-, <u>I</u>RT-like <u>p</u>roteins (ZIP) is well-known for its role in the transport of zinc (Zn), iron (Fe), or other metals across cellular membranes (Eide, 2004). The ZIP family takes its name from the first identified members: <u>z</u>inc <u>r</u>egulated <u>t</u>ransporters (ZRT1 and ZRT2) found in *Saccharomyces cerevisiae* (Zhao & Eide, 1996a, Zhao & Eide, 1996b) and <u>i</u>ron <u>r</u>egulated <u>t</u>ransporter (IRT1) from *Arabidopsis thaliana* (Eide *et al.*, 1996). Since these initial discoveries, ZIP-family proteins have been identified in organisms of various phyla, including 14 in humans (Jeong & Eide, 2013). While ZIP-family proteins principally transport zinc or iron, two human versions, hZIP8 (Park *et al.*, 2015, Fujishiro & Himeno, 2019) and hZIP14 (Scheiber *et al.*, 2019, Aydemir *et al.*, 2017), as well as BmtA from *Borreliella burgdorferi* (Ouyang *et al.*, 2009, Ramsey *et al.*, 2017) primarily transport manganese. Bacterial ZIP proteins fall into the GufA subfamily (Gaither & Eide, 2001), which also contains mammalian members such as hZIP11 (Dempski, 2012, Yu *et al.*, 2013). The first bacterial ZIP protein, ZupT, was identified in *Escherichia coli* (Grass *et al.*, 2002). This initial study proved that it played a role in zinc uptake, and further investigation determined that other metal cations could also be transported by ZupT, albeit with lower affinity (Grass *et al.*, 2005, Taudte & Grass, 2010). Many bacterial species contain putative ZIP-family proteins, but few have been characterized for metal affinity and contribution to growth and virulence.

Here we confirm that ZIP-family proteins should be considered an additional family of bacterial manganese importers. We report that a ZIP-family protein in *S. sanguinis* contributes to manganese uptake in an Δ*ssaACB* mutant, deleted for the ABC-family manganese transporter, and we establish its contribution to IE virulence. We extended this study to additional *S. sanguinis* strains, including those that encode MntH homologs. To our knowledge, we have performed the most extensive analysis of the role of distinct protein families in manganese uptake and virulence that has ever been performed in any *Streptococcus*.

## Results

### Identification of a ZIP-family protein

Given the low concentration of manganese required to improve the growth of the Δ*ssaB* mutant (Crump *et al.*, 2014), we hypothesized that there was a secondary manganese transporter encoded by the *S. sanguinis* SK36 genome. Previously studies in *B. burgdorferi* reported that a ZIP-family protein, BmtA, transported primarily manganese instead of zinc or iron (Ouyang *et al.*, 2009, Ramsey *et al.*, 2017). Since the discovery and characterization of ZupT in *E. coli* (Grass *et al.*, 2002), few bacterial ZIP-family proteins have been characterized for their metal selectivity and only two have been found to contribute to virulence: BmtA (Ouyang *et al.*, 2009) and ZupT in *Clostridioides difficile* (Zackular *et al.*, 2020). *C. difficile* ZupT, like most ZIP-family proteins, was found to principally transport zinc. When examining the genome of SK36, we identified SSA_1413 (SSA_RS06930) as a ZIP-family protein based on comparison to the sequence of BmtA.

### Deletion of the ZIP-family protein gene results in manganese deficiency in an Δ*ssaACB* background

To determine the function of SSA_1413, the knockout strain from an *S. sanguinis* SK36 mutant library (Xu *et al.*, 2011) was utilized. The kanamycin (Kan) resistance cassette and flanking region from the SSX_1413 strain were amplified and transformed into a tetracycline (Tet) resistant Δ*ssaACB* single transport system mutant in an attempt to create a Δ*ssaACB* ΔSSA_1413 double mutant. Initial attempts to introduce the ΔSSA_1413 mutation into the Δ*ssaACB* background were unsuccessful (data not shown). This was not due to lack of competence, as we have previously generated mutants in this strain (Puccio *et al.*, 2020, Murgas *et al.*, 2020). Additionally, this same construct was used to generate the ΔSSA_1413 strain in SK36. Transformants in the Δ*ssaACB* background were obtained only when the brain heart infusion (BHI) agar plates were supplemented with 10 μM Mn^2+^. While the single mutant grew similarly to wild type (WT) in BHI media, the double mutant required 10 μM Mn^2+^ supplementation to grow to WT levels overnight (data not shown). Because of these results and others described below, we concluded that SSA_1413 is a metal transporter, which we have named TmpA for <u>t</u>ransport of <u>m</u>etal <u>p</u>rotein A. This name was also chosen to avoid species- and metal-specific nomenclature.

Serum contains nanomolar concentrations of manganese (Liu *et al.*, 2005) and the oxygen concentration of arterial blood is 12% (Atkuri *et al.*, 2007). In these conditions, growth of WT and Δ*ssaB* mutant strains was determined to be analogous to that observed in our rabbit model of IE, which entails infection of the aortic valve (Crump *et al.*, 2014). At this O_2_ concentration, deletion of *tmpA* from the WT and Δ*ssaACB* backgrounds led to only slight, non-significant differences in growth (Figure 1A). When the O_2_ concentration was reduced to 6% or 1%, growth of the double mutant at 24 h was significantly less than the Δ*ssaACB* parent strain (Figure 1A). In contrast, growth of the single mutant was not statistically different from WT under any of the tested O_2_ concentrations (Figure 1A). Additional studies in BHI showed that again, the Δ*tmpA* mutant grew indistinguishably from WT in all tested O_2_ concentrations, whereas the double mutant grew significantly less than its Δ*ssaACB* parent in all tested O_2_ concentrations (Figure 1B). These results indicate that TmpA contributes to growth in the Δ*ssaACB* background.

**Figure 1.**
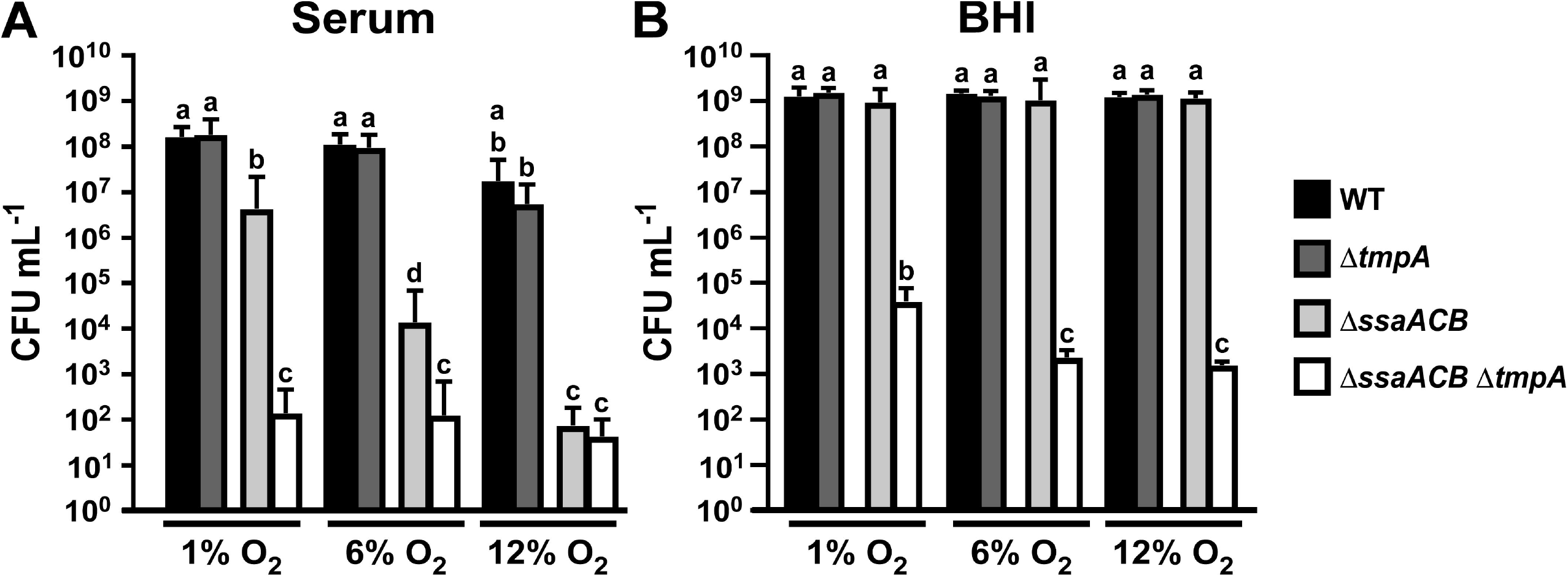
Growth of SK36 manganese-transporter mutants in various O_2_ concentrations. Cultures were grown in (A) rabbit serum or (B) BHI at the given O_2_ concentration for 24 h. Means and standard deviations of at least three independent experiments are displayed. Significant differences between each Δ*tmpA* mutant and its respective parent strain under the same experimental conditions were determined using one-way ANOVA with a Tukey multiple comparisons post-test. Bars with the same letter in each chart are not significantly different from each other (*P* > 0.05).

### Complementation of the Δ*ssaACB* Δ*tmpA* mutant by addition of metals

Although the creation of the Δ*ssaACB* Δ*tmpA* mutant was facilitated by the addition of Mn^2+^ to the media, we wanted to test the effect of addition of Mn^2+^ and other divalent cations on growth of the mutant strains in serum, used here as a biologically relevant manganese-deficient medium. Significant differences in growth of the two strains were observed for every added manganese concentration up to 20 μM (Figure 2). SsaB was also found to transport iron (Crump *et al.*, 2014) and we found that the growth of the Δ*ssaACB* mutant was maximized by the addition of 100 μM Fe^2+^ (Figure 2). Growth of the double mutant was significantly less than the parent in every tested Fe^2+^ concentration. Growth of the two mutant strains was not significantly different in any concentration of added Zn^2+^ (Figure 2), despite the fact that ZIP-family proteins characterized previously primarily transport zinc or iron (Dempski, 2012). These results suggest that TmpA may contribute to both manganese and iron transport in serum.

**Figure 2.**
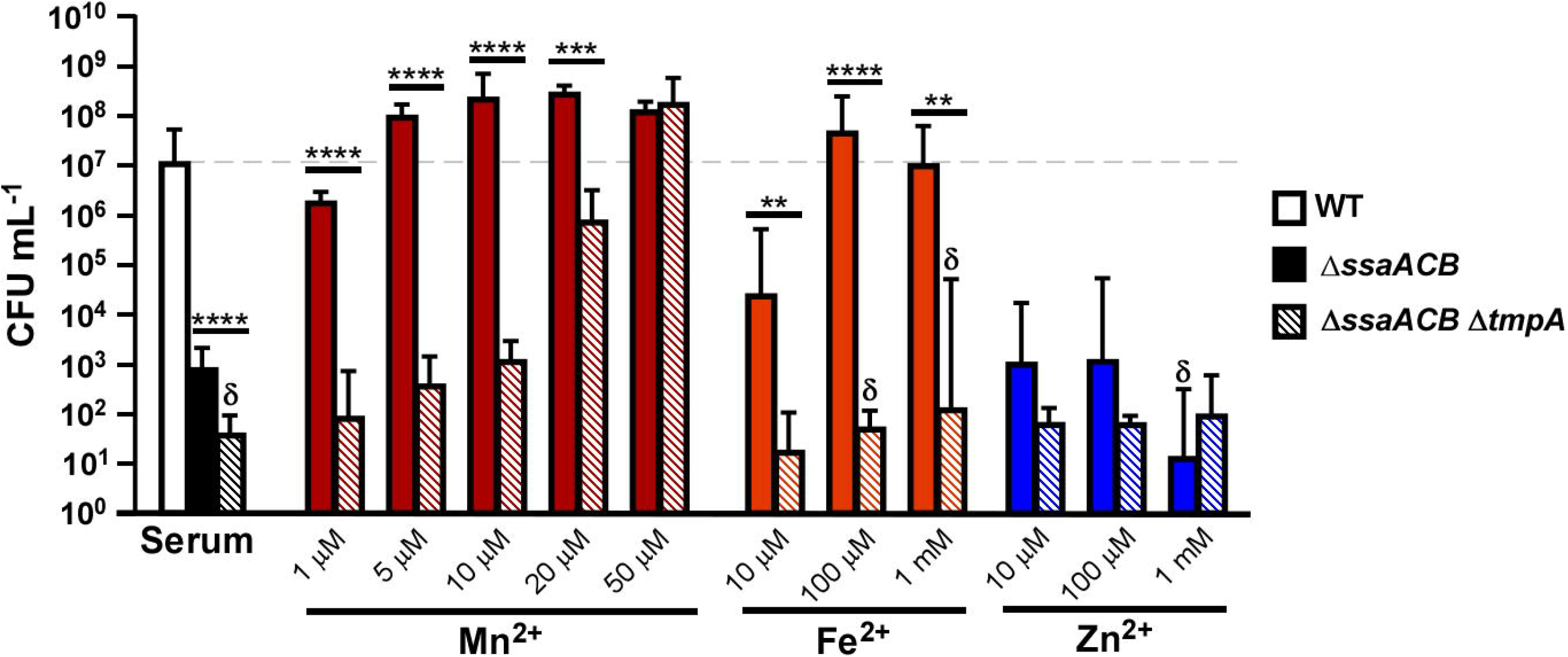
Growth of SK36 Δ*ssaACB* and Δ*ssaACB* Δ*tmpA* mutants in serum with added metals. Cultures were grown in serum at 12% O_2_ with added (A) Mn^2+^, (B) Fe^2+^, or (C) Zn^2+^ for 24 h. Means and standard deviations of at least three independent experiments are displayed. Significant differences between the Δ*ssaACB* parent and Δ*ssaACB* Δ*tmpA* mutant under the same experimental conditions were determined using unpaired two-tailed t-tests (\**P* ≤ 0.05, \*\**P* ≤ 0.01, \*\*\**P* ≤ 0.001, \*\*\*\**P* ≤ 0.0001). Horizontal dashed line indicates WT growth for reference. Bars with δ had at least one replicate that fell below the limit of detection.

In another test of complementation with added metals, 24-h growth of the double mutant on Todd-Hewitt + Yeast Extract (THY) plates required added Mn^2+^ (Figure S1). Neither added Fe^2+^ nor Zn^2+^ had any effect on growth.

### Complementation of the Δ*ssaACB* Δ*tmpA* mutant with inducible expression of *tmpA*

We next sought to determine whether the phenotype we observed in the Δ*ssaACB* Δ*tmpA* mutant could be complemented. To accomplish this, *tmpA* was placed under the control of the *lac* promoter (Phyper-spank) at an ectopic expression site (Turner *et al.*, 2009) in the double mutant strain. When gene expression was induced by addition of 1 mM isopropyl-β-D-thiogalactoside (IPTG), the complemented double mutant grew indistinguishably from WT, surpassing the growth of both the Δ*ssaACB* and the Δ*ssaACB* Δ*tmpA* strains (Figure 3). This result indicates that overexpression of *tmpA* leads to an increase in growth. The addition of 10 μM Mn^2+^ without IPTG to the complemented strain also improved growth to WT levels. Presumably this is due to leakiness of the Phyper-spank promoter, leading to considerable expression of *tmpA* even without IPTG present. We have observed this previously with the same inducible promoter and expression site (Rhodes *et al.*, 2014). These results confirm that the phenotype of the double mutant is due to loss of TmpA rather than any unintended mutation and suggests that even low-level expression of the *tmpA* gene at its native location would be sufficient to augment the growth of the Δ*ssaACB* mutant to the extent indicated in Figure 1. The Δ*ssaACB* mutant was complemented previously (Murgas *et al.*, 2020).

**Figure 3.**
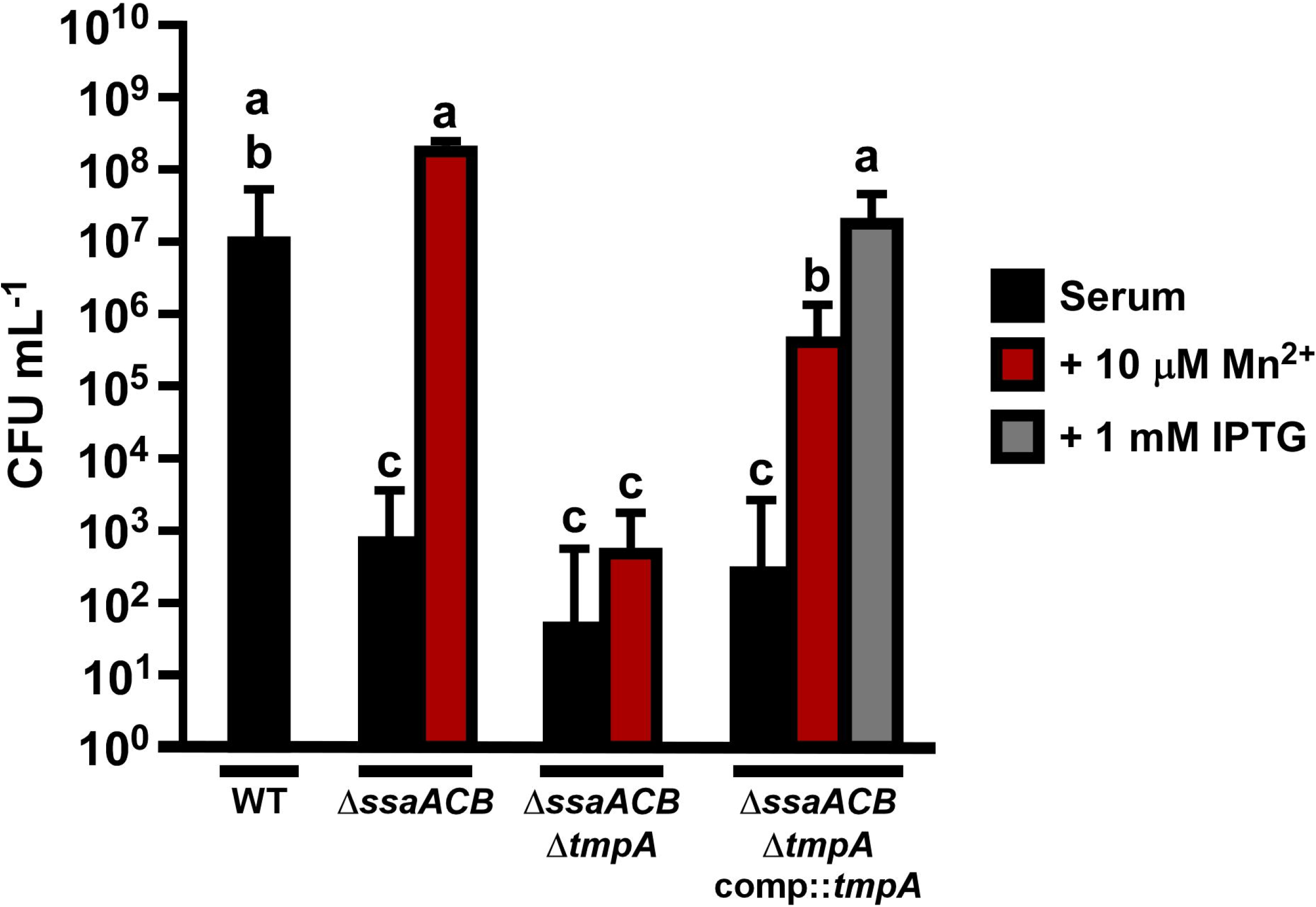
Complementation of serum growth of the Δ*ssaACB* Δ*tmpA* mutant. Cultures were grown in serum at 12% O_2_ for 24 h with 10 μM Mn^2+^ or 1 mM IPTG added as shown. Means and standard deviations of at least three independent experiments are displayed. Significance was determined by a one-way ANOVA with a Tukey multiple comparisons post-test. Horizontal dashed line indicates WT growth for reference. Bars with the same letter in each chart are not significantly different from each other (P > 0.05).

### Assessment of cellular metal content of Δ*tmpA* mutant strains

We then set out to directly test the effect of *tmpA* deletion on cellular metal concentrations. To assess the metal content of each strain, cells were grown overnight then diluted into aerobic (~21% O_2_) BHI with 10 μM of added Mn^2+^, Fe^2+^, or Zn^2+^. The cultures were incubated for several hours, and then cells were collected, washed, digested, and analyzed by inductively coupled plasma optical emission spectroscopy (ICP-OES). There was no significant difference between either Δ*tmpA* mutant and its respective parent strain in plain BHI for any metal tested (Figure 4), including magnesium (Figure S2A). With added Mn^2+^, the double mutant imported significantly less manganese than the Δ*ssaACB* parent strain (Figure 4A). This trend was not observed for iron or zinc when 10 μM of each respective metal was added (Figure 4B-C). Additionally, there were no difference in magnesium, manganese, iron, or zinc when a different metal was added (Figure S2).

**Figure 4.**
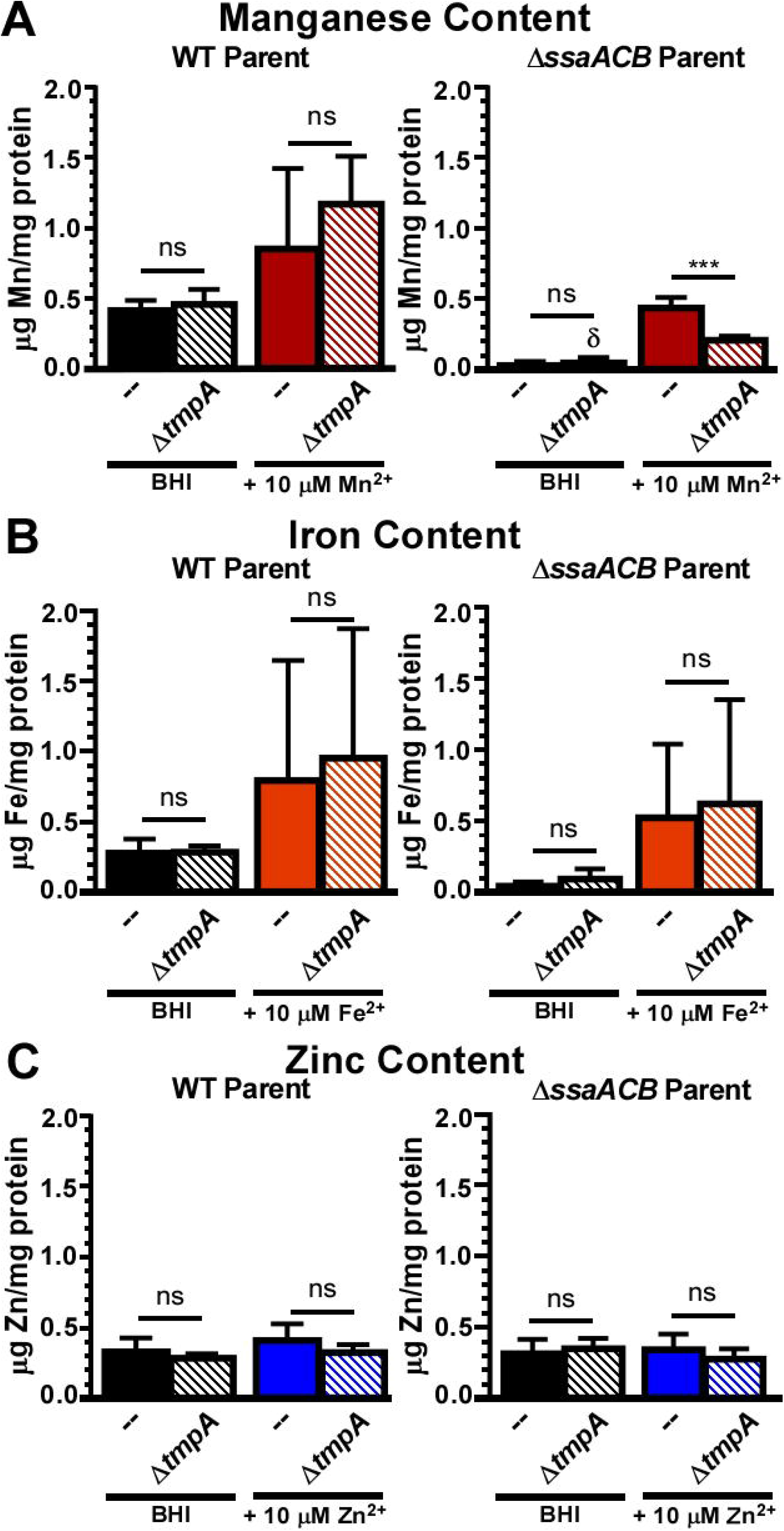
Metal content of SK36 manganese-transporter mutants in BHI. Metal content of cells grown in BHI ± 10 μM Mn^2+^ (A), Fe^2+^ (B), or Zn^2+^ (C) in atmospheric O_2_ (~21%). Metal concentration was measured by ICP-OES and normalized to protein concentration. Means and standard deviations of at least three independent experiments are displayed. Significance was determined by one-way ANOVA with Bonferonni’s multiple comparisons test comparing each mutant and its respective parent in each growth condition. Bars with δ indicate that at least one value fell below the lowest standard.

Since the manganese-dependent phenotype of *tmpA* was only observed in the Δ*ssaACB* background, this left the possibility that a zinc-transport phenotype was masked by the presence of high-affinity zinc transporters. To test this possibility, we first attempted to create a Δ*adcCBA* mutant, deleted for the three adjacent genes encoding the zinc ABC transporter AdcCBA (Dintilhac *et al.*, 1997) but were unsuccessful in obtaining transformants (data not shown). This result indicated that the *adcCBA* operon may be essential for growth. Thus, we decided to generate an Δ*adcC* single gene mutant using the knockout construct from Xu *et al.* (2011), since in this study, individual mutants of all three components were made successfully, suggesting that deletion of each component separately was tolerated even if deletion of the entire operon was not. As expected, this mutant grew normally in BHI (data not shown) and exhibited poor growth in the presence of the metal chelator TPEN when grown in Chelex-treated BHI (cBHI) (Figure S3A). TPEN is often used to reduce available zinc (Ganguly *et al.*, 2021). An Δ*adcC* Δ*tmpA* strain was then created to assess whether TmpA transports zinc in addition to manganese. Growth of the Δ*adcC* Δ*tmpA* strain was not statistically different from the Δ*adcC* strain in zinc-depleted conditions (Figure S3A). We then evaluated whether there was a difference between the growth of the Δ*adcC* and Δ*adcC* Δ*tmpA* mutants when various concentrations of Zn^2+^ were added to the media and found that there were no significant differences in any added Zn^2+^ concentration (Figure S3B). We then measured the cellular metal content of these mutant strains in cBHI (Figure S3C). There was no significant difference in zinc levels between either Δ*tmpA* mutant and its respective parent, although the slight decrease in the zinc content of the Δ*adcC* Δ*tmpA* strain relative to its parent when 10 μM Zn^2+^ was added suggests that TmpA may make a minor contribution to zinc transport under these conditions.

### Expression of the *tmpA* gene under various metal and oxygen concentrations

Bacterial metal transporters are often negatively regulated by the metals that they transport, in most cases by binding of the metal to a protein that represses transcription of the transporter gene(s) (Johnston *et al.*, 2006, Kehres & Maguire, 2003). Therefore, as another approach to investigate the metal specificity of TmpA, we investigated the effect of various metals on the regulation of its gene. WT and Δ*ssaACB* cells were grown in BHI and then incubated with 100 μM of either Mn^2+^, Fe^2+^, Zn^2+^, or EDTA (Figure 5A). A BHI-only sample was used as the control for comparison. Unfortunately, expression of *tmpA* was not significantly affected by the addition of any tested metal nor by the depletion of manganese by EDTA (Figure 5A). The MntR ortholog in *S. sanguinis*, SsaR, has been shown to negatively regulate the expression of *ssaB* in the presence of added manganese (Crump *et al.*, 2014). Thus, the *ssaB* gene and the *aphA-3* (Kan resistance) gene that replaced the *ssaACB* genes in the Δ*ssaACB* mutant were included as positive controls for manganese-dependent regulation and as expected, expression significantly decreased in the presence of added manganese (Figure 5A).

**Figure 5.**
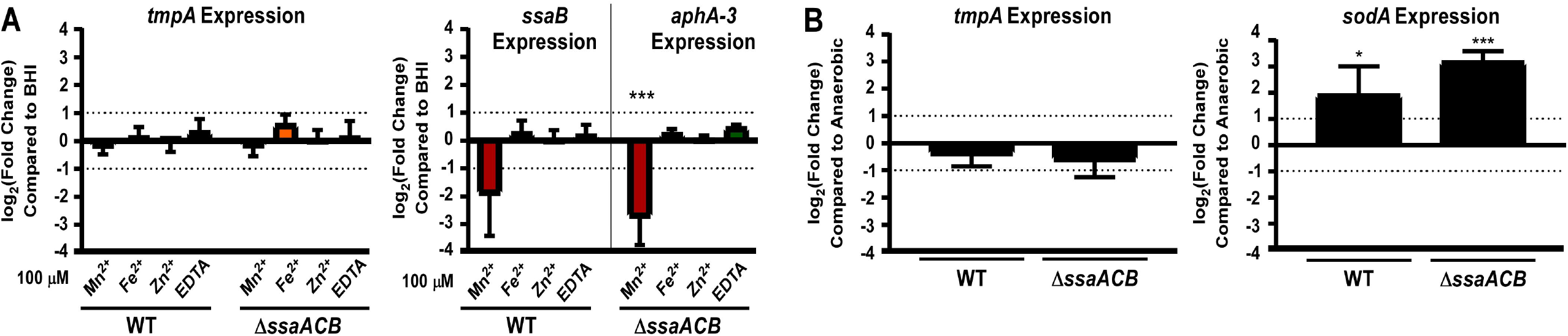
Expression of *tmpA* in SK36 WT and Δ*ssaACB* cells when exposed to various metal concentrations and O_2_ conditions. (A) Relative transcript levels of cells exposed to 100 μM Mn^2+^, Fe^2+^, Zn^2+^, or EDTA for 15 min as compared to BHI alone. (B) Relative transcript levels of cells exposed to oxygen after anaerobic growth. Means and standard deviations of three independent experiments are displayed. Significant differences between the ΔCt values for the experimental condition compared to the control condition within the same strain were determined by unpaired two-tailed t test. Only conditions with |log_2_(fold change) values| > 1 were tested for significance. \**P* ≤ 0.05, \*\*\**P* ≤ 0.001

Given that manganese is important in oxidative environments (Waters, 2020), we then assessed whether expression of *tmpA* was affected by O_2_ concentration by measuring transcript levels before and after exposure to oxygen. Expression of *tmpA* did not change significantly from anaerobic to aerobic conditions (Figure 5B). The expression of the positive control *sodA* significantly increased after oxygen exposure, as expected (Crump *et al.*, 2014). Thus, we have yet to discover a condition that leads to differential expression of *tmpA* and suspect that it may be constitutively expressed. We attempted to assess production of the TmpA protein in *S. sanguinis* by western blot but were unsuccessful (data not shown).

### Contribution of TmpA to virulence in a rabbit model of infective endocarditis

Given the severe reduction in endocarditis virulence of the Δ*ssaB* (Crump *et al.*, 2014) and Δ*ssaACB* (Baker et al., 2019) mutants, we decided to test whether TmpA also contributes to virulence. We employed a rabbit model in which a catheter was inserted through the carotid artery to induce minor aortic valve damage (Turner *et al.*, 2009). Bacterial strains were then introduced into the bloodstream by co-inoculation into a peripheral ear vein. Infected vegetations were recovered the following day from euthanized animals and homogenized. The recovered bacteria were enumerated by dilution plating on selective antibiotics (Turner *et al.*, 2009). Recovery of the Δ*tmpA* mutant was not significantly different from WT and both were recovered in significantly higher numbers than the Δ*ssaACB* strain, which was only recovered in one of six rabbits (Figure 6A). These results indicate that in a WT background, TmpA is not required for virulence in our model, likely because SsaACB can import sufficient manganese to support growth from the low levels found in blood.

**Figure 6.**
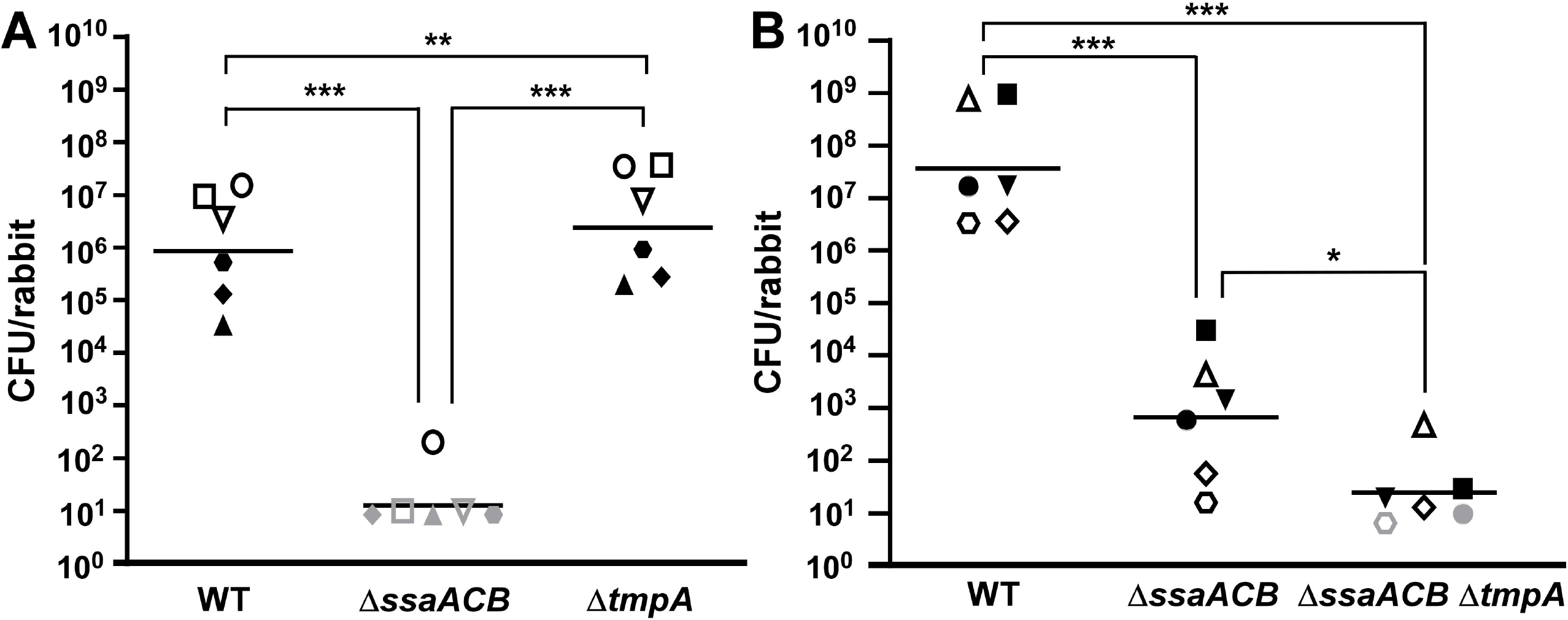
Virulence of SK36 manganese-transporter mutants in a rabbit model of IE. Rabbits were co-inoculated with the marked WT strain, the Δ*ssaACB* mutant, and either (A) the Δ*tmpA* or (B) the Δ*ssaACB* Δ*tmpA* mutant strain. In (A), the inoculum sizes for the three strains were approximately equal and normalized to inocula for each experiment. In (B), the inoculum for WT was 20 times lower than the inoculum of each Δ*ssaACB* mutant, thus final recovery was multiplied by 20 to normalize to the inocula of the other two strains. Within each panel, like symbols indicate bacteria of each strain recovered from the same animal (n =6 over two independent experiments). Females are represented by open symbols and males are represented by closed symbols. Gray symbols indicate recovery was below the limit of detection. Horizontal lines indicate geometric means. \**P* ≤ 0.05, \*\**P* ≤ 0.01, \*\*\**P* ≤ 0.001 indicate significant differences using repeated measures ANOVA with a Tukey multiple comparisons test.

We next wanted to assess the contribution of *tmpA* to virulence in a Δ*ssaACB* background; however, the exceedingly low recovery of the Δ*ssaACB* mutant made it unlikely that a further reduction in virulence would be detectable in our model. Therefore, for the next experiment, the WT inoculum level was decreased two-fold while the Δ*ssaACB* and Δ*ssaACB* Δ*tmpA* mutant inocula were increased 10-fold, resulting in a 20-fold difference relative to WT. The Δ*ssaACB* strain was recovered from all six rabbits but at a significantly lower level than WT (Figure 6B). The recovery of the Δ*ssaACB* Δ*tmpA* mutant was significantly lower than the Δ*ssaACB* mutant. These results suggest that the loss of TmpA in the Δ*ssaACB* background resulted in a further decrease in virulence, indicating that it may be playing a secondary role in manganese uptake that is only evident when the primary manganese transporter is absent or inactive.

### Contribution of specific residues of TmpA to its function

To examine the differences between TmpA and other ZIP-family proteins, we aligned the amino acid sequence of TmpA to those of two metal-selective ZIP transporters: ZIPB (zinc) and BmtA (manganese) (Figure S4). We then used Protter (Omasits *et al.*, 2014) to generate a 2D depiction of the protein within a membrane based on the transmembrane domains (TMDs) predicted from the alignment (Figure 7). All ZIP-family proteins characterized thus far have been integral membrane proteins with eight TMDs and are typically predicted to have both C- and N-termini facing extracellularly (Guerinot, 2000). They usually contain two canonical motifs: (*i*) a variable length (Hx)_n_ motif in the cytoplasmic loop between TMD III and IV (Eide, 2004) and (*ii*) a conserved HNxPEG motif in TMD IV (Lin *et al.*, 2010). As with BmtA and ZIPB, TmpA has several histidine residues in the variable loop region between TMDs III and IV but only two follow the (Hx)_n_ pattern. Both protein sequences contain the conserved HNxPEG motif in TMD IV. From the alignment (Figure S4), we found four putative metal-binding residues – E67, N173, E240, and N251 – that were different from the confirmed metal-binding residues from the crystal structure of ZIPB (Zhang *et al.*, 2017).

**Figure 7.**
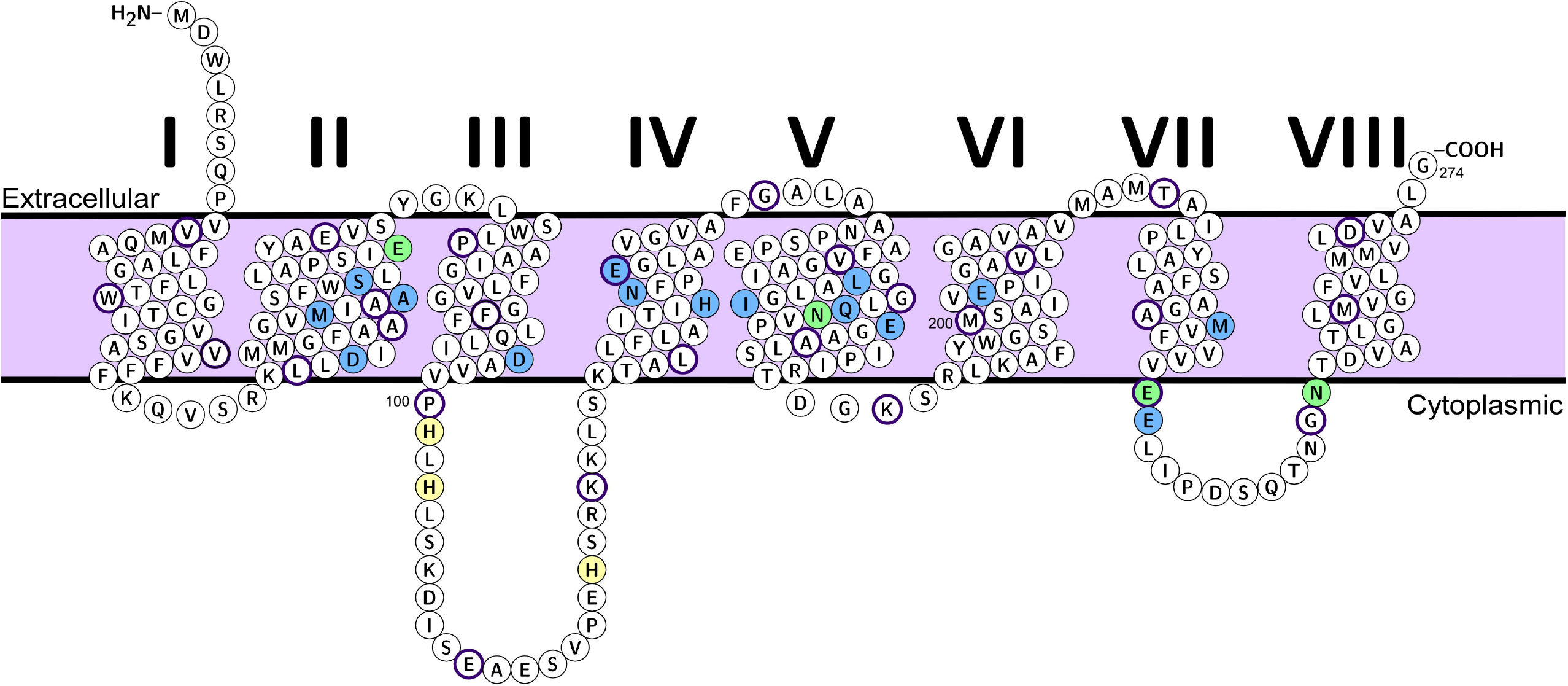
Depiction of predicted TmpA tertiary structure within a membrane. TmpA amino acid sequence depicted in a cell membrane using Protter (Omasits *et al.*, 2014). TMDs of TmpA were predicted based on alignment to the crystal structure of ZIPB (Figure S4). Every 10^th^ residue is circled in dark purple. Amino acids in blue and green are conserved putative metal-binding residues based on the crystal structure of ZIPB bound to zinc (Zhang *et al.*, 2017). Those in green are metal-binding residues that are not conserved between ZIPB and TmpA. Yellow residues are histidines thought to contribute to metal transport (Zhang *et al.*, 2019) but have not been confirmed by the crystal structure.

Residue E67 was found near the extracellular side of the protein, while N173 was predicted be within the central region of the protein (Figure 7) in the “M2” site of the binuclear center (Zhang *et al.*, 2017). Residues E240 and N251 were located in the cytoplasmic loop between TMDs 7 and 8 (Figure 7). Because of the disordered nature of long loops, the main loop between TMDs 3 and 4 was not crystallized; thus, no metal-binding residues were identified in that region of the crystal structure (Zhang *et al.*, 2017). Three of the four residues in TmpA matched those in BmtA (Figure S4). We decided to mutate all four of these residues in TmpA to alanines to determine the contribution of each side chain to the function of the protein (Morrision & Weiss, 2001). We also mutated these four residues to the corresponding residue from ZIPB to determine whether this would affect the function. Since we could not determine a phenotype for the Δ*tmpA* single mutant, we made these site-directed mutants (SDM) in the Δ*ssaACB* background.

We then assessed the growth of these mutants in comparison to WT, the Δ*ssaACB* mutant, and the Δ*ssaACB* Δ*tmpA* mutant in rabbit serum at 6% O_2_ (Figure 8A). All E67 and E240 mutants grew similarly to the Δ*ssaACB* parent strain, indicating that these residues are not essential for TmpA protein function. N251H also grew indistinguishably from the parent strain but N251A grew significantly worse, which suggests that this residue may be important for transport, but that histidine is also capable of performing the same function. This was not unexpected, as histidine is known to coordinate manganese (Martin & Giedroc, 2016) and the metal-binding residue at position 251 varied between TmpA and BmtA (Figure S4). Both N173 mutants grew poorly, which suggests that this residue is critical for function.

**Figure 8.**
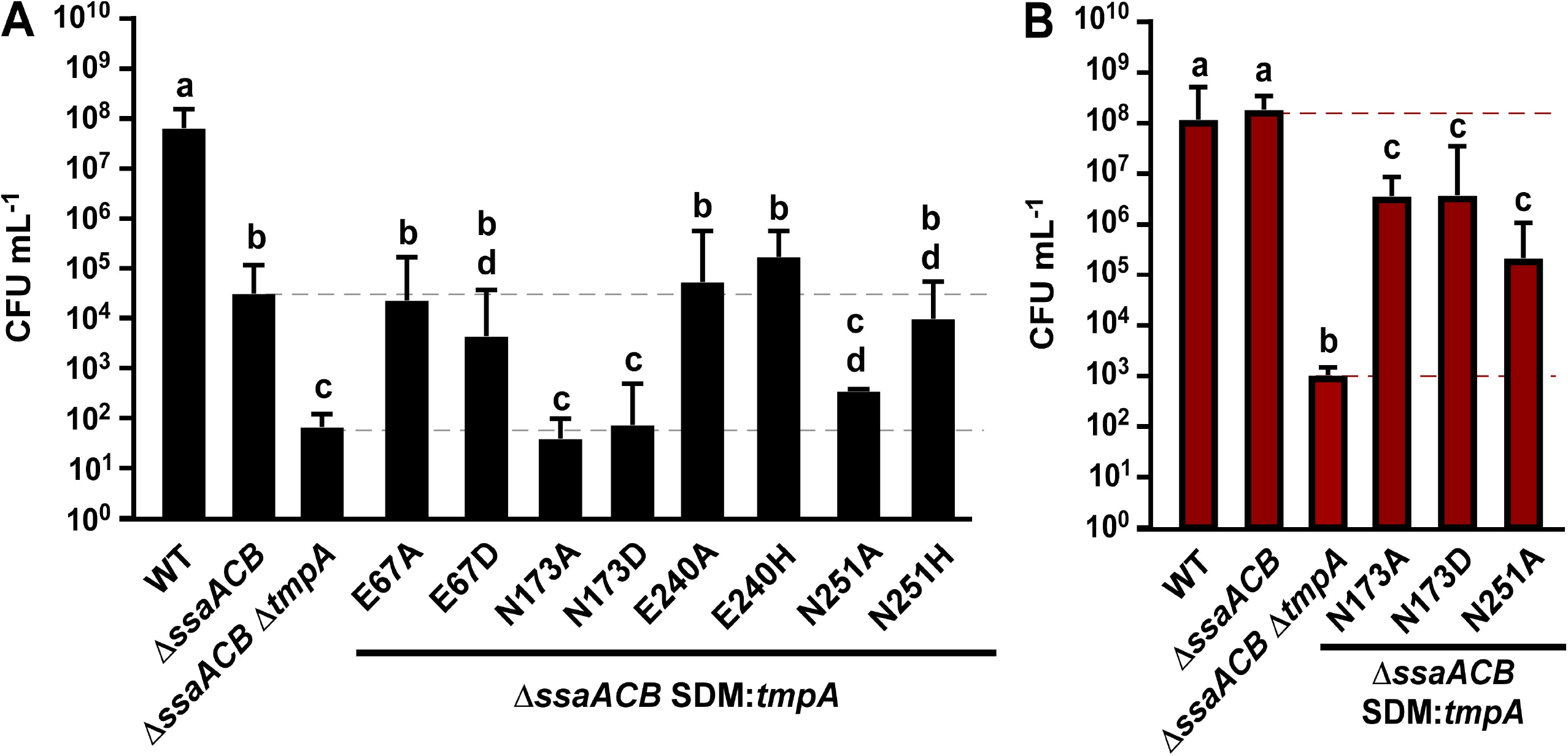
Growth of Δ*ssaACB* SDM: *tmpA* mutants. Growth of single amino acid (site-directed mutagenesis; SDM) mutant versions of TmpA in the Δ*ssaACB* background was assessed in serum at 6% O_2_. (B) Mutants deficient in growth as compared to the Δ*ssaACB* parent at 6% O_2_ in A were evaluated for growth under the same conditions + 5 μM Mn^2+^. Means and standard deviations of at least three independent experiments are shown. Significance was determined by one-way ANOVA with a Tukey multiple comparisons test. Bars with the same letter are not significantly different from each other (*P* > 0.05).

Despite our best efforts, we were unable to detect TmpA by western blot (Puccio, 2020). However, we were able to conclude that tagged protein was still being produced, as strains with the Strep-Tag® II tag fused to TmpA at two different sites grew similarly to the parent strain when 5 μM Mn^2+^ was added (Figure S5). The clear exception was the N-terminally tagged version, which suggests that the N-terminus may be important either for function or for localization to the membrane. Additionally, we attempted to overexpress the TmpA protein in *E. coli* strains optimized for heterologous membrane protein expression. While we were able to confirm the presence of TmpA by western blot (Puccio, 2020), we were unsuccessful in obtaining sufficient protein quantities to perform cell-free transport assays in liposomes, which would have been ideal for measuring the contribution of each residue to transport.

We thus considered two extreme explanations for the failure of the N173A, N173D, and N251A mutants to display growth that was significantly better than the Δ*ssaACB* Δ*tmpA* mutant in Figure 8AFigure 8A: (*i*) the N173 and N251 mutations interfered with the secretion, localization, or stability of TmpA; or (*ii*) the SDM were normal with regard to these properties, and their lack of detectable activity was due to the importance of these residues for manganese transport. To distinguish between these two possibilities, the three mutants that grew poorly in serum in 6% O_2_ were assessed for growth after addition of 5 μM Mn^2+^ (Figure 8B). When Mn^2+^ was added, the Δ*ssaACB* parent grew to WT-like levels, whereas the Δ*ssaACB* Δ*tmpA* mutant’s growth was still significantly lower than WT. Each of the mutants grew to a level that was lower than the Δ*ssaACB* parent but higher than the Δ*ssaACB* Δ*tmpA* mutant. The results for all three SDM are in agreement with the second model, in which the lack of activity in unamended serum is due to reduced metal transport rather than loss of the protein.

### Model of TmpA based on the ZIPB crystal structure

To determine whether the two N173 mutations would be expected to severely affect metal transport function, we modeled TmpA using the crystal structure of ZIPB as a template (Figure S6A). The full-length proteins share 40% identity, while the crystal structure of ZIPB (PBD: 5TSA) shares 34% identity with TmpA due to lack of structural data for several loops. Additionally, TMD III in TmpA was shorter than that of ZIPB, and thus was depicted missing one of the helical loops in the model (Figure S6A). The reason for this short TMD is unclear, as the length of all other TMDs appear to match well.

We then modelled the N173D mutation in TmpA (Figure S6B). The other residues that constitute the putative M2 binding site moved to accommodate the negative charge of the aspartic acid, which resulted in a change in the size and shape of the M2 site. The change in shape and size is, of course, speculative, as proteins *in vivo* are flexible and may be able to accommodate changes such as these. Nevertheless, the predicted structural changes, in conjunction with the obvious change in charge, could well explain the severity of this mutation’s effect on metal transport and growth observed in Figure 8.

The position of the proteins within the cellular membrane was then predicted using Orientation of Proteins in Membranes (https://opm.phar.umich.edu/). As described in Zhang *et al.* (2017), the protein has been crystallized with a tilt (Figure S7A). The model of TmpA fits well in the predicted membrane with this tilt (Figure S7B).

### Evaluation of serum growth of manganese-transporter mutants in additional *S. sanguinis* strains

Given the considerable genotypic and phenotypic heterogeneity that we previously observed between different strains of *S. sanguinis* (Baker *et al.*, 2019), we wanted to determine whether our findings concerning manganese transport in SK36 would apply to other strains as well. We therefore examined the manganese transporters in four additional *S. sanguinis* strains: SK49, SK408, SK678, and VMC66. Like SK36, SK49 is an oral isolate. By ICP-OES, we found previously that it accumulated less manganese when cultured in BHI than 14 of the 17 strains examined (Baker *et al.*, 2019). VMC66 (Kitten *et al.*, 2012), SK408, and SK678 were all isolated from the blood of endocarditis patients and ranked first, second, and third, respectively, in manganese levels in the same assay (Baker *et al.*, 2019).

Manganese transporter mutants—Δ*ssaACB*, Δ*tmpA*, and Δ*ssaACB* Δ*tmpA*—were generated for each strain. These mutants and their parent strains were then assessed for serum growth at 6% O_2_ (Figure 9). As with SK36 (Figure 1A), all Δ*tmpA* mutant strains were indistinguishable from WT and all Δ*ssaACB* strains were deficient for growth. Interestingly, the Δ*ssaACB* Δ*tmpA* mutants of SK678 and VMC66 grew similarly to their respective Δ*ssaACB* parent strains, whereas the SK49 and SK408 versions grew to significantly lower densities than their parent strains. However, it is apparent that in SK678 and VMC66, the *ssaACB* deletion produced a greater defect on growth than in the other backgrounds (Figure 9C-D).

**Figure 9.**
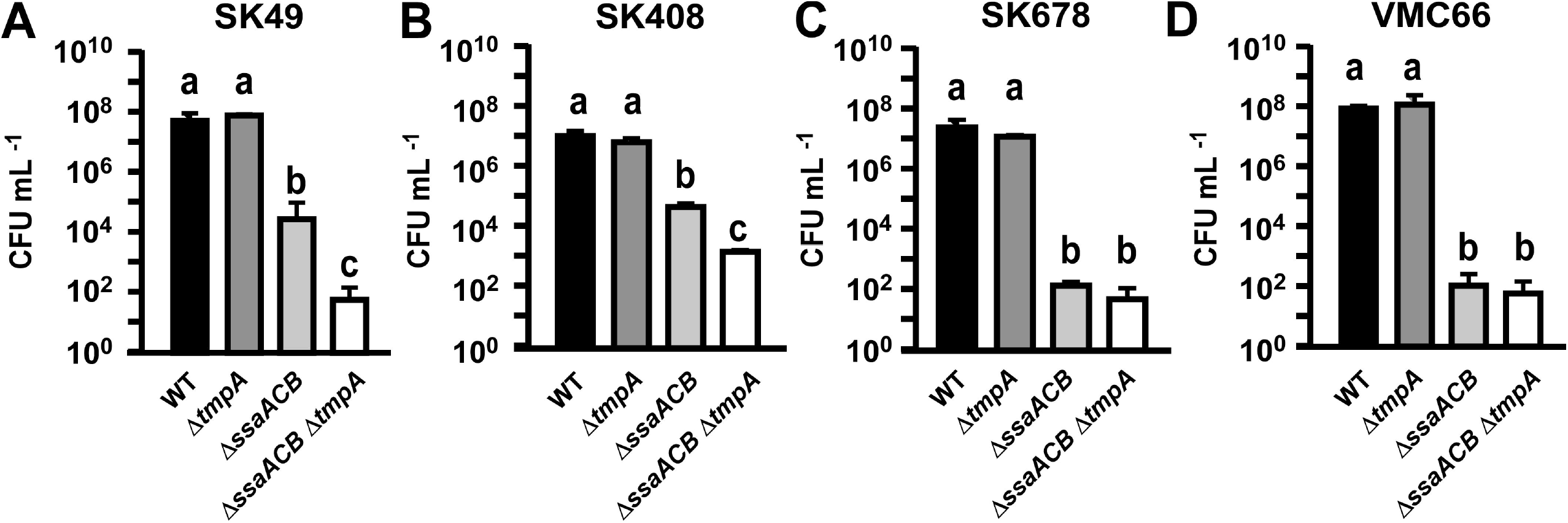
Growth of manganese-transporter mutants of other *S. sanguinis* strains. Growth of each strain and its respective mutants in serum at 6% O_2_ was assessed by plating on BHI agar at 24 h. Means and standard deviations of at least three independent experiments are displayed. Significance was assessed by one-way ANOVA with a Tukey multiple comparisons test. Bars that share a letter within a chart are not significantly different from each other (*P* > 0.05).

To determine if the poor growth of the Δ*ssaACB* parents prevented us from detecting an additional effect of *tmpA* deletion, growth of the SK678 and VMC66 groups was assessed in 1% O_2_ (Figure S8). Under these conditions, we observed a significant difference between each Δ*ssaACB* Δ*tmpA* mutant and its Δ*ssaACB* parent. In summary, the mutant phenotypes observed in the SK36 background were similar to those of mutants in other strains of *S. sanguinis*.

### Relative contribution of TmpA and MntH to growth and virulence of *S. sanguinis* VMC66

Of interest to this study, at least eight *S. sanguinis* strains with genome sequences available in GenBank encode gene orthologous to the NRAMP protein MntH found in *S. mutans* (Kajfasz *et al.*, 2020) and other gram-positive bacteria (Kehl-Fie *et al.*, 2013). NRAMP proteins are known to import manganese (Bozzi *et al.*, 2016, Ehrnstorfer *et al.*, 2014) and contribute to endocarditis virulence in *Enterococcus faecalis* (Colomer-Winter *et al.*, 2018). It was unexpected then that the VMC66 Δ*ssaACB* mutant performed so poorly in the 6% O_2_ serum growth study (Figure 9D). To determine whether these NRAMP proteins may contribute to manganese uptake and endocarditis virulence in *S. sanguinis*, knockout mutants were generated in the VMC66 WT and Δ*ssaACB* strains and serum growth was assessed. At 6% O_2_, the Δ*mntH* strain grew to a significantly lower level than the WT parent but higher than the Δ*ssaACB* mutant (Figure 10A). Both double mutant strains grew similarly to their Δ*ssaACB* parent (Figure 10A).

**Figure 10.**
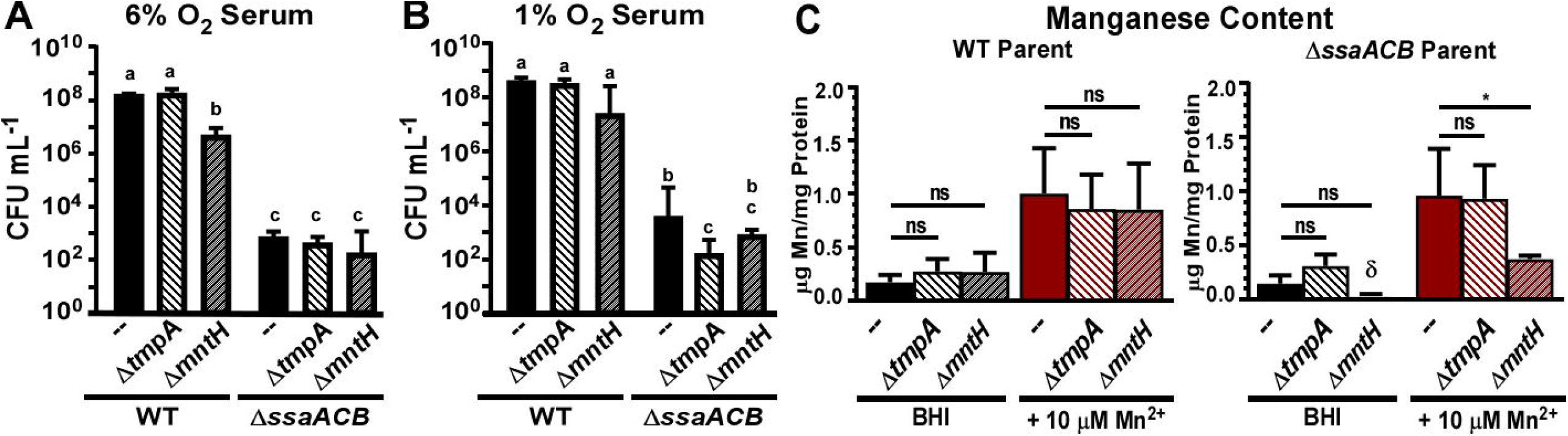
Growth and manganese content of VMC66 manganese-transporter mutants. Growth of VMC66 and its respective manganese-transporter mutants in serum at (A) 6% O_2_ and 1% O_2_ was assessed by plating on BHI agar after 24 h. Means and standard deviations of at least three independent experiments are displayed. Significance was assessed by one-way ANOVA with a Tukey multiple comparisons test for T_0_ and T_24_ separately. T_24_ bars that share a letter within a chart are not significantly different from each other (*P* > 0.05). T_0_ bars with § or & are significantly different from each other (*P* ≤ 0.05). (C) Manganese content of each strain in BHI ± 10 μM Mn^2+^ was measured by ICP-OES and normalized to protein concentration. Means and standard deviations of at least three independent experiments are displayed. Significance was determined by one-way ANOVA with Dunnett’s multiple comparisons tests for each secondary transporter mutant as compared to the parent strain under the same conditions (\**P* ≤ 0.05). The δ indicates that at least one replicate was below the lowest standard.

These results indicate that MntH can contribute to growth even in cells possessing the other two transporters, but its contribution is much less than that of SsaACB. As seen in Figure 9, the drastic growth reduction in the Δ*ssaACB* mutant masked the contribution of TmpA in this background at 6% O_2_ and this could have been true in this experiment for MntH as well. Thus, we decided to assess growth at 1% O_2_ (Figure 10B). When we lowered the oxygen concentration, we once again observed that the Δ*ssaACB* Δ*tmpA* mutant of VMC66 grew significantly worse than the Δ*ssaACB* parent. Both Δ*mntH* and Δ*ssaACB* Δ*mntH* grew slightly but not significantly less than their respective parent strains.

We then assessed the relative contribution of each secondary transporter to manganese import by measuring cellular metal content by ICP-OES (Figure 10C). Similar to our previous experiments with SK36 manganese transporter mutants, the low cellular manganese levels that were observed for the VMC66 Δ*ssaACB* mutants grown in BHI alone made it difficult to evaluate potential differences. To circumvent this issue, we also measured manganese content when 10 μM Mn^2+^ was added to the BHI. We found that there were no significant differences between the manganese content of either Δ*tmpA* mutant and its respective parent strain under either condition (Figure 10C). The same was true of other tested metals: iron, zinc, and magnesium (Figure S9). However, when Mn^2+^ was added, manganese levels in the Δ*ssaACB* Δ*mntH* mutant were significantly lower than in the Δ*ssaACB* parent, suggesting that MntH contributes more to manganese transport in this background.

To determine the relative contribution of each manganese transporter to VMC66 virulence, WT, Δ*tmpA*, Δ*ssaACB, and* Δ*mntH* strains were tested in our rabbit model of IE (Figure 11A). To selectively plate the WT, we utilized a VMC66 Spc^R^ WT strain generated previously (Baker *et al.*, 2019). The Δ*mntH* mutant was recovered at similar levels to this marked WT strain. The only strain to be recovered at significantly lower levels than WT was the Δ*ssaACB* strain, highlighting its importance as the primary manganese transporter in multiple *S. sanguinis* strains.

**Figure 11.**
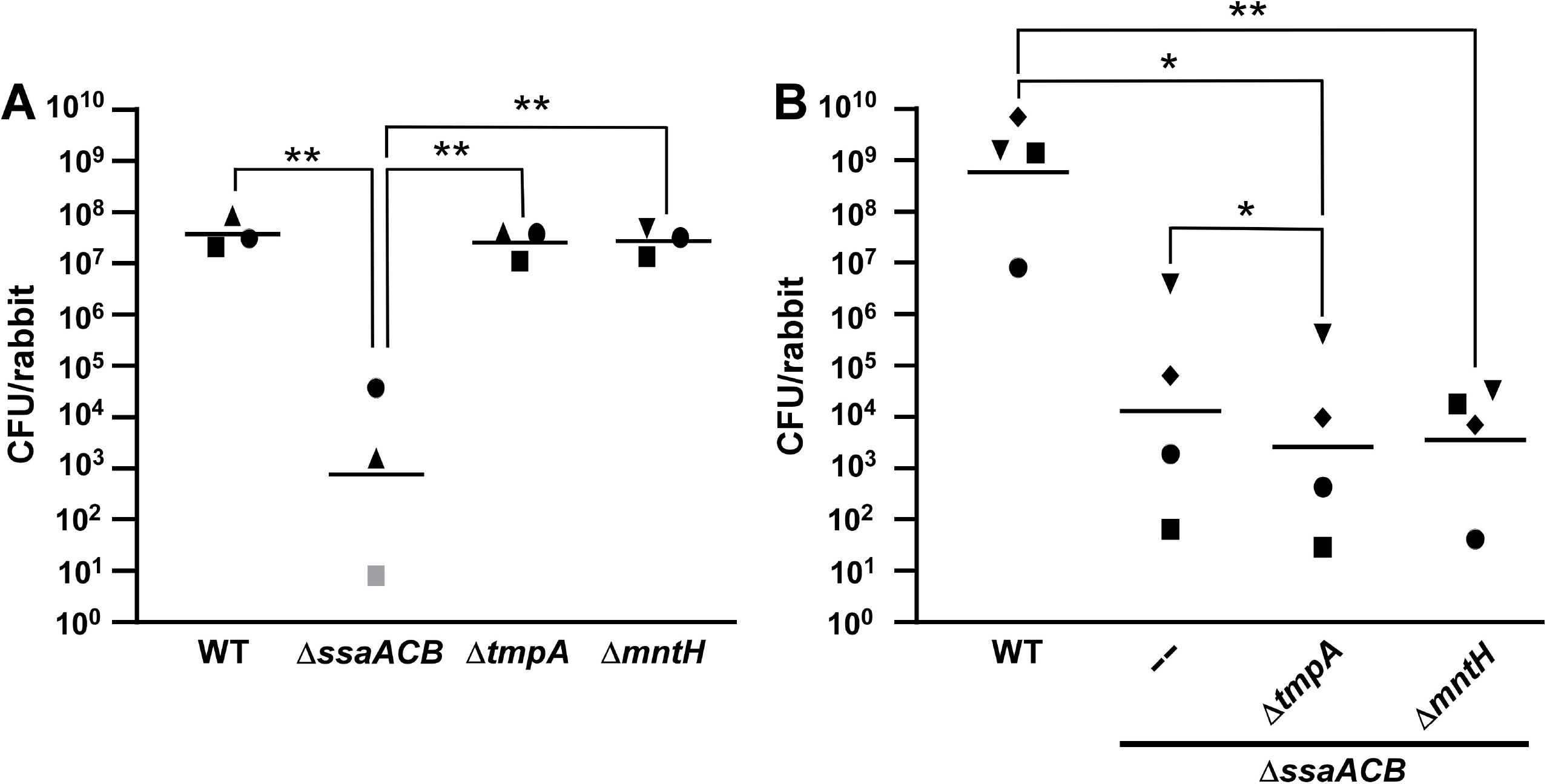
Virulence of VMC66 manganese-transporter mutants in a rabbit model of IE. Rabbits were co-inoculated with the marked WT strain, the Δ*ssaACB* mutant strain, and either (A) the Δ*tmpA* and Δ*mntH* single mutant strains or (B) the Δ*ssaACB* double mutant strains. In (A), the inoculum sizes for the three strains were approximately equal and normalized to inocula for each experiment. In (B), the inoculum for WT was 20 times lower than the inoculum of each Δ*ssaACB* mutant, thus final recovery was multiplied by 20 to normalize to the inocula of the other two strains. Within each panel, symbols of the same shape indicate bacteria of each strain recovered from the same animal, where (A) n = 3 and (B) n = 4. All rabbits were male. Gray symbols indicate recovery was below the limit of detection. Geometric means are indicated by horizontal lines. \**P* ≤ 0.05, \*\**P* ≤ 0.01 indicate significantly different from other strains using repeated measures ANOVA with a Tukey multiple comparisons test.

To test the relative contribution of each secondary transporter to virulence, the double mutants were also tested in our IE rabbit model (Figure 11B). To ensure sufficient recovery of the Δ*ssaACB* mutants, we once again increased the inoculum size of each of these mutants and decreased the inoculum size of WT so that the WT inoculum was twenty times less than that of the three mutants. We were able to recover colonies of every strain from each rabbit and we saw a significant difference between the WT and both double mutant strains. However, the difference between WT and the Δ*ssaACB* mutant fell short of significance (*P*-value = 0.0589). We observed a significant difference between the Δ*ssaACB* Δ*tmpA* mutant and its parent but neither was significantly different from the Δ*ssaACB* Δ*mntH* mutant. This suggests that TmpA may contribute to virulence, even when MntH is present.

### Phylogenetic analysis of TmpA homologs

We next wanted to assess the phylogenetic distribution of the TmpA protein in streptococci. BLASTP (Altschu *et al.*, 1990) was used to search the non-redundant protein sequence database at NCBI with the amino acid sequence of TmpA. A TmpA homolog was present in many streptococcal species for which at least one complete genome sequence was present. Exceptions included *S. pneumoniae*, *Streptococcus mitis*, *Streptococcus oralis*, *Streptococcus pyogenes*, and *Streptococcus vestibularis*, all of which appear to lack a TmpA homolog or contain only partial sequences. Additionally, a phylogenetic analysis showed that previously characterized ZIP proteins from other genera of bacteria, such as BmtA and ZIPB, were more closely related to the main group of streptococcal ZIP proteins than were the homologs from *S. mutans* and *Streptococcus ratti* (Figure S10A). The TmpA homolog of *Streptococcus sobrinus,* a member of the “Mutans” group of streptococci (Nobbs *et al.*, 2009), also clustered with the main group of streptococcal proteins rather than with those from *S. mutans* and *S. ratti*. ZupT proteins from *E. coli* and *C. difficile* were more similar to the *S. mutans* protein than to TmpA from *S. sanguinis*. Furthermore, we included the hu man ZIP-family proteins hZIP11 and hZIP8 with the intention of using them as outgroups. We found that hZIP11 was more closely related to most of the bacterial ZIP-family proteins than were the those of *S. mutans* and *S. ratti*, whereas the hZIP8 protein functioned as an actual outgroup. These results, along with the observation that our ZIP protein phylogenetic tree looks very different from a 16S rRNA-based tree (Nobbs *et al.*, 2009), indicate that genes encoding ZIP-family proteins were likely incorporated into the genome of each species or various progenitors at different times, which has been observed previously in streptococci, as opposed to being derived from a common ancestor (Richards *et al.*, 2014). The GC content of the *S. sanguinis* genome is 43.40% (Xu *et al.*, 2007) whereas *tmpA* is 50.42%, indicating that it could have been acquired by horizontal gene transfer (Ravenhall *et al.*, 2015). The GC content of the *S. mutans* UA159 genome is 36.82% (Ajdić *et al.*, 2002) and its ZIP protein SMU_2069 (here designated ZupT_Sm_) is 37.1%, which suggests that it could have evolved with *S. mutans* or been acquired from another organism with a similar GC content.

To investigate this further, we evaluated the phylogeny of SsaB orthologs in streptococci (Figure S10B). Most species contained orthologs of SsaB, including those that were lacking a ZIP-family protein, such as *S. mitis*, *S. oralis*, *S. pyogenes*, and *S. vestibularis.* Interestingly, *Streptococcus suis* and *S. ratti* lack an SsaB ortholog. In this case, *S. mutans* again groups separately from the rest of the streptococci (100% of 500 replicate trees inferred by bootstrap analysis), including another “Mutans” group *Streptococcus*, *S. downei*, but the *S. mutans* ortholog (SloC) clusters more closely with the other streptococcal proteins than do representatives from more distantly related species, including *Staphylococcus aureus* and *B. bronchiseptica*. We then looked closer at the *S. mutans* ZIP protein, ZupT_Sm_. When we performed a BLASTP search against GenBank with ZupT_Sm_ as the query, most of the hits were either other Firmicutes or streptococcal species that are closely related to *S. mutans* (Figure S11). The *S. sanguinis* TmpA sequence was included as an outgroup and we included the same sequence of ZupT from *C. difficile* as in Figure S10A for reference. Additionally, when examining the top non-streptococcal hits by BLAST search for each ZIP-family protein, we noted that the top hits to TmpA after those from the first group of firmicutes, most were from γ-proteobacteria. When ZupT_Sm_ was used as the query, almost all of the top hits were firmicutes of different genera from those identified in the TmpA search. These analyses further support independent incorporation of TmpA/ZupT orthologs in *S. mutans* and the other streptococci at different times in history, although we can now add that they are also likely from different sources.

### Sequencing of the SK36 Δ*ssaACB* Δ*tmpA* double mutant

After completing the experiments in this manuscript, we sequenced the whole genomes of the Δ*ssaACB* mutant (JFP173) and the Δ*ssaACB* Δ*tmpA* mutant (JFP227). Unfortunately, we discovered the presence of a nonsense mutation in SSA_1414 (W139*) in JFP227. SSA_1414 is annotated as MutT, an 8-oxo-dGTP diphosphatase. We confirmed this mutation by Sanger sequencing and found that it was unique to this strain, as it was not present in the Δ*tmpA* single mutant (JFP226), which was made with the same PCR product as JFP227. We recreated markerless versions of this mutation in the WT and Δ*ssaACB* backgrounds and found no significant differences in serum growth for either version (Figure S12). We next generated a clean Δ*ssaACB* Δ*tmpA* strain and confirmed by Sanger sequencing that the SSA_1414 gene was intact. We compared the growth of this new mutant (JFP377) to that of JFP227 (Figure S13A-C) under various conditions. There was no significant difference in growth between these strains. We then measured the metal content of cells grown in BHI with 10 μM Mn^2+^ using ICP-OES and observed no significant difference in any metal examined (Figure S13D).

## Discussion

With this study, it has now been confirmed that at least two different bacterial species, *S. sanguinis* and *B. burgdorferi*, utilize a ZIP-family protein for manganese uptake. This firmly establishes the ZIP family as an additional bacterial manganese importer family. Additionally, we confirm that an NRAMP-family protein, MntH, contributes to manganese uptake in at least one *S. sanguinis* strain in which it is naturally encoded. These findings confirm the existence of secondary manganese transporters in *S. sanguinis* (Figure 12). Despite the fact that there was an unintended mutation in the Δ*ssaACB* Δ*tmpA* mutant in a nearby gene, SSA_1414, we were able to demonstrate that a clean mutant shares the same phenotype (Figure S12) and that the Δ*ssaACB* Δ*tmpA* mutant could be complemented by expression of the *tmpA* gene at a different chromosomal site (Figure 3).

**Figure 12.**
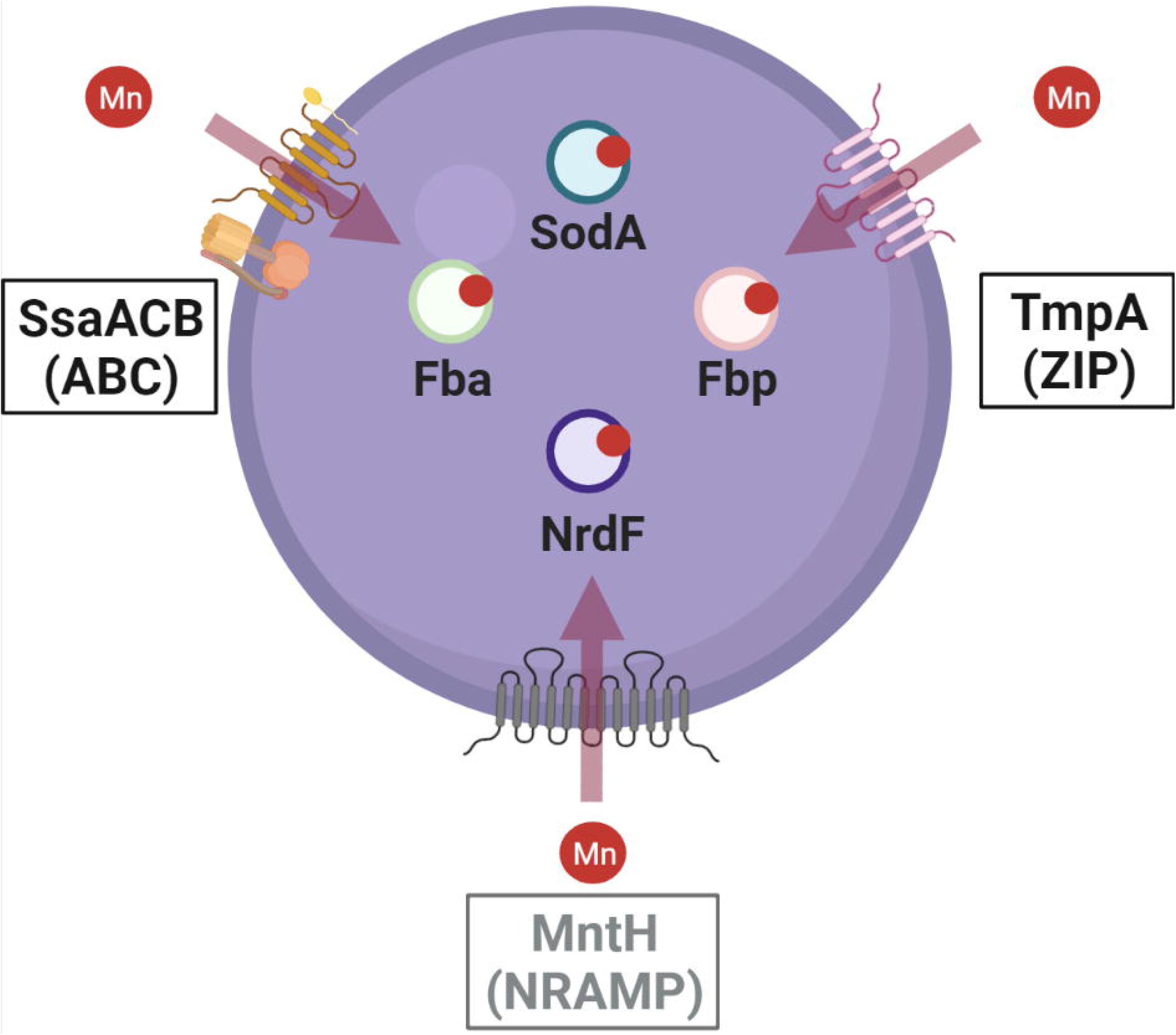
Summary of known manganese transporters and manganese-utilizing enzymes in *S. sanguinis*. Representation of a single *S. sanguinis* cell with confirmed and predicted manganese-transporting or -utilizing proteins. Red circles represent manganese ions. Arrows indicate direction of transport across the cellular membrane. Manganese-dependent enzymes are depicted as circles within the cell. Transporters are depicted with putative TMDs and other components, such as the ATPase and lipoprotein components of SsaACB. MntH (gray) has only been found in eight sequenced strains of *S. sanguinis.* Figure was generated with Biorender.com

We showed that the Δ*ssaACB* mutant accumulates less iron and manganese, in agreement with our previous results (Crump *et al.*, 2014, Murgas *et al.*, 2020). The findings that low levels of manganese restore serum growth to the Δ*ssaACB* mutant, that higher manganese levels are required for restoration of growth to the Δ*ssaACB* Δ*tmpA* mutant, and that less cellular manganese is detected in the Δ*ssaACB* Δ*tmpA* mutant compared to its Δ*ssaACB* parent are all consistent with the conclusion that TmpA transports manganese and that this activity is important in the background of an Δ*ssaACB* mutant. The finding that supplemental iron restores serum growth to the Δ*ssaACB* mutant has not been reported previously, although in our earlier paper we did not attempt complementation with more than 10 μM supplemental iron, and we reported that it did not reproducibly restore growth (Crump *et al.*, 2014), which is not inconsistent with our findings from this study. It thus appears that supplemental iron is capable of restoring serum growth, although the levels required are perhaps ~20-fold greater than required for manganese. We presume that mononuclear iron is substituting for manganese as a cofactor in enzymes that can use either metal (Anjem & Imlay, 2012).

The effects of the Δ*tmpA* mutation in relation to iron are less easily understood than the effects on manganese. We found that, as with manganese, the deletion of *tmpA* from the Δ*ssaACB* mutant resulted in iron being less effective at restoring serum growth; indeed, while ~20-fold more manganese was required to restore growth to the double mutant relative to its parent, no amount of added iron (up to 1 mM) was sufficient to restore growth. This result could suggest that TmpA is a secondary transporter of iron as well as manganese and that unlike manganese, which we presume is transported into the double mutant by one or more tertiary transporters, there is no tertiary transporter for iron under these conditions. Yet, unlike with manganese, the Δ*ssaACB* Δ*tmpA* double mutant and its Δ*ssaACB* parent were indistinguishable with regard to iron uptake in the presence or absence of supplemental iron (Figure 4), which is inconsistent with TmpA-mediated iron transport.

We suggest that these seemingly inconsistent findings can be explained by hypothesizing that there are certain core functions that require manganese, and no amount of iron can restore growth if manganese levels are insufficient to meet these needs. Therefore, the failure of iron to complement the double mutant is due to the loss of TmpA-mediated manganese transport rather than TmpA-mediated iron transport. If we presume that increased levels of manganese are needed for the core functions as levels of oxygen and, therefore, oxygen-dependent reactive oxygen species such as superoxide and hydrogen peroxide increase (Anjem & Imlay, 2012), this would also be consistent with the data in Figure 1 showing that as oxygen levels decrease, the Δ*ssaACB* mutant is capable of ever more growth in serum. It should also be noted that particularly in blood if not also in the oral cavity, free iron is present in vanishingly low levels (Diaz-Ochoa *et al.*, 2014); thus, the significance of iron transport by the SsaACB transporter is not clear.

Despite the fact that most ZIP-family proteins transport zinc, there were no significant differences between any Δ*tmpA* mutant and its respective parent in either zinc levels or zinc-deficient growth. Any difference in zinc levels between the Δ*adcC* and Δ*adcC* Δ*tmpA* mutants with 10 μM added Zn^2+^ was insignificant and given that this concentration is likely biologically unattainable, the combined data suggest that zinc transport, if any, is minimal.

We have been unable to find a differential phenotype for the Δ*tmpA* mutant as compared to WT. In our accompanying manuscript (Puccio *et al.*, 2021b), we tested its growth in low-pH BHI and saw no significant differences. We also assessed the growth in a biofilm assay in competition with *S. mutans* observed no difference (data not shown). These results, along with those described in this study, strongly suggest that its function is secondary to SsaACB.

The reason that most organisms encode multiple transporters for each metal is still contested, although it highlights the importance of these transition metals. The presence of multiple transporters with varying affinity could be due to the transport systems functioning optimally under different environmental conditions or it could protect from the loss of function of one of the transporters (Raimunda *et al.*, 2011, Eide, 2012, Dempski, 2012). For example, excess zinc was found to inhibit the manganese transport function of the SsaB ortholog in *S. pneumoniae*, PsaA (Counago et al., 2014). If this is also true for SsaB, TmpA’s function could become essential under the same condition. In our accompanying manuscript (Puccio *et al.*, 2021b), we found that loss of Δ*ssaACB* in SK36 was detrimental to growth in low-pH media, which we hypothesize is due to acid sensitivity of TmpA. In VMC66, lack of Δ*ssaACB* had no effect in the same conditions, likely because MntH is still functional at the pH tested. Further studies will be required to confirm the identity and selectivity of other metal transporters in *S. sanguinis*.

In VMC66, the only single manganese transport system mutant that showed significantly decreased virulence was the Δ*ssaACB* mutant (Figure 11A). The Δ*ssaACB* Δ*tmpA* mutant was recovered at significantly lower levels than the Δ*ssaACB* mutant whereas the Δ*ssaACB* Δ*mntH* mutant was not. However, when the two double mutants were compared to each other, they were not significantly different (Figure 11B). Thus, we cannot confidently conclude that the Δ*tmpA* mutation had a greater effect on the virulence of the Δ*ssaACB* mutant than the Δ*mntH* mutation did.

Expression of *tmpA* was not significantly affected by addition of metals or by depletion with EDTA. EDTA is a non-specific metal chelator which was chosen to represent metal-deplete conditions. While EDTA is not specific for any metal, it has an affinity for manganese (log_β1_ of 14.1 and 24.8 for Mn^2+^ and Mn^3+^, respectively, as compared to 16.7 for Zn^2+^ and 8.7 for Mg^2+^) (Perrin & Dempsey, 1974). When added to WT or Δ*ssaACB* cultures growing in a fermentor, ICP-OES analysis revealed that its primary effect on cellular metal levels was to reduce the concentration of manganese (Puccio *et al.*, 2020). Thus, if *tmpA* were regulated by manganese depletion, we would have expected to see a change in expression after the addition of EDTA.

While our transcriptional analysis is not exhaustive, it is plausible that *tmpA* expression is constitutive in *S. sanguinis*. In *E. coli*, the gene encoding ZupT is constitutively transcribed at low levels (Grass *et al.*, 2005). In our RNA-seq analysis of EDTA-treated cells, expression of *tmpA* decreased slightly, yet significantly at T_50_ but initial expression at T_−20_ was low, as were other genes in the operon (Puccio *et al.*, 2020). This result may have been an artifact of data normalization, because in our study examining the transcriptome after pH reduction, expression of *tmpA* remained low and constant in both WT and Δ*ssaACB* strains, despite a decrease in cellular manganese levels (Puccio *et al.*, 2021).

In a recent study, Zhang *et al.* (2020) confirmed that in the human hZIP4, the M1 site is essential for metal transport and the M2 site facilitates optimal transport activity. Here we report that the modification of N173 to A or D in the predicted M2 site of TmpA resulted in the reduced function of the protein as a manganese transporter (Figure 8). This indicates TmpA cannot function efficiently without an asparagine in this position, despite the similar size and features of the aspartic acid. These results suggest that the charge of the side chain at this position in TmpA likely contributes to either transport function, metal selectivity, or both. Future studies using cell-free metal uptake assays and structural analysis will be required to confirm our results.

The discovery that streptococci have different lineages of ZIP-family proteins suggests that these secondary transporters may be valuable to some species. However, this protein family is not essential in all streptococci as some species, such as the human pathogen *S. pyogenes* and the oral commensals *S. oralis* and *S. mitis*, have been evolutionarily successful without a ZIP-family protein. In some instances, MntH is encoded in the genomes of these species but *S. pyogenes* has only two sequenced strains that encode a MntH homolog, suggesting that the other strains either rely entirely on the primary ABC manganese transporter or encode an unidentified secondary transporter. At the other extreme, some streptococci encode all three transporters, similar to *S. sanguinis* VMC66 examined here. *S. mutans* is an example; however, the contribution of ZupT_Sm_ to manganese transport has yet to be examined. A recent study found that it did not contribute significantly to zinc transport (Ganguly *et al.*, 2021). Another study from the same group utilizing *S. mutans* manganese-transport mutants deleted for *sloC,* the *ssaB* homolog, and *mntH* found that the inactivation of MntH alone did not produce any obvious phenotype, whereas the Δ*sloC* and Δ*sloC* Δ*mntH* mutants were severely deficient under manganese-restricted conditions (Kajfasz *et al.*, 2020). Further studies in both *S. mutans* and *S. sanguinis* would improve the understanding of the relative contribution of each of these proteins to growth, metal transport, and virulence and possibly reveal more about the origin of the distinct lineages.

In conclusion, we discovered that this ZIP-family protein, TmpA, contributes to manganese uptake and virulence in several strains of *S. sanguinis,* which is evident only after the deletion of the primary transporter. These results lay the foundation for future studies of manganese-transporter inhibitors for the prevention of IE, as both transporters could be targeted simultaneously to prevent the generation of spontaneous mutants that could subvert single-target treatment. Indeed, a study targeting BmtA to prevent *B. burgdorferi* virulence showed promising results (Wagh *et al.*, 2015), although this study was completed before the crystallization of any ZIP-family protein. Additionally, inhibitors of SsaB orthologs have been under investigation, as they could potentially be used to treat other streptococcal diseases in addition to IE (Obaidullah *et al.*, 2018, Bajaj *et al.*, 2015). Furthermore, given that MntH does not significantly contribute to virulence but remains a functional manganese transporter at low pH (Puccio *et al.*, 2021b), its presence could lessen the effect of anti-SsaB drugs on the oral microbiota and may influence selection of potential probiotic anti-caries streptococcal strains. Although future studies will be required, we hypothesize that any anti-SsaB drugs would likely remain effective at preventing IE without affecting growth or competitiveness of any MntH-encoding streptococcal strain in the oral cavity. Taken together, these studies enhance our understanding of manganese transporters in *S. sanguinis* and the importance of manganese for the growth and virulence of this opportunistic pathogen.

## Materials and Methods

### Bacterial strains and growth conditions

The *S. sanguinis* strains used in this study are listed in **Error! Reference source not found.**. Primers and plasmids used to generate the mutant strains are listed in **Error! Reference source not found.** & S3, respectively. *S. sanguinis* strains SK36, SK49, SK408, and SK678 are human isolates from Mogens Killian, Aarhus University, Denmark, characterized for virulence previously (Baker *et al.*, 2019). VMC66 is a human blood isolate from Virginia Commonwealth University Medical Center Hospital (Kitten *et al.*, 2012, Baker *et al.*, 2019). All strains were grown in overnight cultures from single-use aliquots of cryopreserved cells, diluted 1000-fold in BHI media (Beckinson Dickinson). Mutant strains were incubated with the appropriate antibiotics: kanamycin (Kan; Sigma-Aldrich) at 500 ug mL^−1^; tetracycline (Tet; Sigma-Aldrich) 5 μg mL^−1^; erythromycin (Erm; Sigma-Aldrich) at 10 μg mL^−1^; chloramphenicol (Cm; Fisher Scientific) at 5 μg mL^−1^; and spectinomycin (Spc; Sigma-Aldrich) at 200 μg mL^−1^. The cultures were then incubated at 37°C for 16-20 h with the atmospheric condition set to 1% (1% O_2_, 9.5% H_2_, 9.5% CO_2_ and 80% N_2_), 6% (6% O_2_, 7% H_2_, 7% CO_2_ and 80% N_2_), or 12% (12% O_2_, 4 % H_2_, 4% CO_2_ and 80% N_2_) oxygen using a programmable Anoxomat™ Mark II jar-filling system (AIG, Inc.). Overnight cultures of SK36 Δ*ssaACB* Δ*tmpA* mutants and the VMC66 Δ*ssaACB* Δ*mntH* strain were grown with 10 μM MnSO_4_ added.

Gene knockout mutant constructs were either generated previously (Xu *et al.*, 2011) or by gene splicing by overlap extension (SOEing) PCR (Ho *et al.*, 1989) where the gene(s) of interest were replaced with an antibiotic resistance gene or cassette. Transformations were performed using the protocol described previously (Paik *et al.*, 2005). Briefly, an overnight culture of the parent strain was grown in Todd Hewitt (Beckinson Dickinson) broth with horse serum (Invitrogen), then diluted 200-fold and incubated at 37°C. Optical density (OD_600_) of tube cultures was determined using a ThermoScientific BioMate 3S UV-VIS spectrophotometer. Knockout construct DNA (100 ng) and *S. sanguinis* competence stimulating peptide (70 ng) were added to the culture (OD_600_ ~0.07) and incubated at 37°C for 1.5 h prior to selective plating on BHI agar plates with antibiotics at concentrations listed above. All plates were incubated for at least 24 h at 37°C under anaerobic conditions in an Anoxomat jar with a palladium catalyst. The SK36 Δ*tmpA* strains were derived from SK36 WT and Δ*ssaACB*::*tetM* (JFP173), where *tmpA* was replaced with *aphA-3* using the SSX_1413 strain and primers from the SK36 knockout mutant library (Xu *et al.*, 2011). The double mutant version was grown on BHI plates with 10 μM MnSO_4_ added.

Markerless mutants were generated using a mutation system described previously (Cheng *et al.*, 2018, Xie *et al.*, 2011). Briefly, the in-frame deletion cassette (IFDC) was amplified from the *S. sanguinis* IFDC2 strain and combined with flanking region from *tmpA* using gene SOEing. The parent strains were then transformed as described above, plating on BHI agar plates containing Erm. A gene SOEing product merging the two sides of *tmpA* with the desired nucleotide changes was then generated. This SOEing product was then used to transform the Erm^R^ colonies from the first transformation. Immediately prior to plating on agar plates containing 20 mM 4-chloro-phenylalanine (4-CP; Sigma-Aldrich), the cells were washed twice with phosphate buffered saline (PBS) to remove remaining media. The presence of the desired mutation was confirmed by Sanger sequencing. This method was also used to add the Strep-Tag II® sequence to each of the tagged mutants in its native genome location.

The complemented *tmpA* strain was generated using gene SOEing, which placed the *tmpA* gene under control of the Phyper-spank promoter (Rhodes *et al.*, 2014) and downstream of the Spc resistance cassette, *aad9.* This construct was inserted into the SSA_0169 gene on the chromosome (Turner *et al.*, 2009). The SSA_0169::*tmpA*::ST (Loop III) strain was generated using same method, with the Strep-Tag II® sequence added between residues 109 and 110 of TmpA.

JFP36 is a previously generated marked WT strain (Turner *et al.*, 2009) used for the *in vivo* experiments. All experiments with the SK36 version of the Δ*ssaACB* mutant used the Tet^R^ version (JFP173) with the following exceptions: the qRT-PCR experiment (Figure 5; Kan^R^; JFP169) and one animal experiment (Figure 6B; Tet^R^ Spec^R^; JFP234). Due to conflicts with antibiotic resistance, strain JFP234 was generated by amplifying the *aad9* gene and flanking DNA from the SSA_0169 locus in strain JFP56 (Turner *et al.*, 2009) and introducing this product into JFP173, thus generating a Δ*ssaACB* mutant that is Spc^R^ to allow for selective plating in rabbit experiments. This strain was assessed for growth in serum and was indistinguishable from JFP173 (data not shown).

The Δ*ssaACB,* Δ*tmpA,* and Δ*mntH* mutants in strains other than SK36 were generated by using gene SOEing with primers specific to each strain background when identical primers were not possible. Since the SSA_0169 ectopic expression site from strain SK36 described above is not present in most other backgrounds examined, we previously generated a Spc^R^ VMC66 WT strain with the *aad9* gene inserted into a new, highly conserved ectopic expression site (Baker *et al.*, 2019). Due to issues with overlapping antibiotic selection markers, another Δ*ssaACB* Δ*tmpA* mutant strain was generated for use exclusively in the rabbit virulence study that was Cm^R^ and Erm^R^. Given that both Δ*ssaACB* Δ*tmpA* and Δ*ssaACB* Δ*mntH* were Cm^R^, they were plated on Erm and Tet plates, respectively.

### Growth studies

Overnight BHI pre-cultures of each strain were made as described above. Tubes containing either 100% pooled rabbit serum (Gibco), BHI, or Chelex-treated (BioRad) BHI supplemented with 1 mM CaCl_2_ and 1 mM MgSO_4_ (cBHI) were pre-incubated at 37°C. The tubes were placed in Anoxomat jars set to either 1% O_2_, 6% O_2_, or 12% O_2_ (12% O_2_, 4.3% CO_2_, 4.3% H_2_). Plating was used to enumerate colony forming units (CFUs). To determine CFUs, samples were sonicated for 90 s using an ultrasonic homogenizer (Biologics, Inc) to disrupt chains prior to dilution in PBS and plated using an Eddy Jet 2 spiral plater (Neutec Group, Inc.).

For experiments in which metal was added, we employed the Puratronic™ line of metals (Alfa Aesar; MnSO_4_·H_2_O, FeSO_4_·H_2_O, and ZnSO_4_·H_2_O; 99.999% guaranteed purity). Each solution was made in Chelex-treated deionized water (cdH_2_O) and added to the serum tubes immediately prior to inoculation. Fe^2+^ stocks were made fresh immediately prior to each experiment. For growth assessment of the complemented mutant, 1 mM IPTG (Fisher Scientific) was added to the serum tubes immediately prior to inoculation. For the Δ*adcC* mutant experiments, 1 μM TPEN (Sigma-Aldrich) and various concentrations of Zn^2+^ were added to the cBHI immediately prior to inoculation.

### Spot plating

Todd-Hewitt (BD) with 1% Yeast Extract (BD) agar plates were made with 100 μM Puratronic™ metals (MnSO_4_, FeSO_4_, or ZnSO_4_). Overnight pre-cultures were grown as described above and then serially diluted in PBS. Cultures (10 μL) were spotted onto the plates aerobically, allowed to dry, and then incubated at 0% O_2_ overnight before imaging.

### Metal analysis

Overnight BHI pre-cultures were grown as described above. Two tubes containing 38 mL BHI or cBHI were pre-incubated at 37°C for each experimental condition. The following day, 3 mL of the overnight culture was used to inoculate each 38-mL media tube. For cBHI experiments, pre-cultures were centrifuged and resuspended in warm cBHI prior to inoculation. Puratronic™ metals were prepared as described above and added immediately prior to inoculation. Inoculated cultures were placed back in the incubator. After several hours of growth, cells were harvested by centrifugation at 3,740 x *g* for 10 min at 4°C. The supernatant was decanted and the cell pellet was washed twice with cold cPBS (PBS treated with Chelex-100 resin for 2 h, then filter sterilized and supplemented with EDTA (Invitrogen) to 1 mM). The pellet was then divided for subsequent acid digestion or protein concentration determination. Trace metal grade (TMG) nitric acid (15%; Fisher Chemical) was added to one portion of the pellet. The pellet was digested using an Anton Paar microwave digestion system and a modified Organic B protocol: 120°C for 10 min, 180°C for 20 min, with the maximum temperature set to 180°C. The digested samples were then diluted 3-fold with cdH_2_O. Metal concentrations were determined using an Agilent 5110 inductively coupled plasma-optical emission spectrometer. Concentrations were determined by comparison with a standard curve created with a 10 μg ml^−1^ multi-element standard (CMS-5; Inorganic Ventures) diluted in 5% TMG nitric acid. Pb (Inorganic Ventures) was used as an internal standard (10 μg ml^−1^). The other portion of the pellet was resuspended in PBS and mechanically lysed using a FastPrep-24 instrument with Lysing Matrix B tubes (MP Biomedicals) as described previously (Rhodes *et al.*, 2014). Insoluble material was removed by centrifugation. Protein concentrations were determined using a bicinchoninic acid (BCA) Protein Assay Kit (Pierce) as recommended by the manufacturer, with bovine serum albumin as the standard. Absorbance was measured in a black-walled, flat-bottom 96-well plate (Greiner) using a microplate reader (BioTek).

### Quantitative real time polymerase chain reaction

Overnight cultures of SK36 WT and Δ*ssaACB* (JFP169) strains were grown as described above. They were then diluted 10-fold into BHI and incubated aerobically (~21% O_2_). Once cells reached mid-log phase (OD_600_ ~0.6), 6 mL of culture was separated into tubes for each condition and 100 μM Puratronic™ metal (MnSO_4_, ZnSO_4_, or FeSO_4_) or EDTA (Invitrogen) was added. A culture tube with no additives was included as the control. Tubes were incubated aerobically without a jar at 37°C for 15 min. To collect cells, the tubes were swirled in a dry ice/ethanol bath for 30 s prior to centrifugation for 10 min at 3,740 x *g* at 4°C. The supernatant was discarded and the samples stored at −80°C. In some experiments, overnight cultures of SK36 and JFP169 were grown in an anaerobic chamber (Coy Laboratory Products). Cells were then diluted 10-fold into BHI pre-incubated anaerobically at 37°C. At mid-log phase, cultures were separated and 6 mL of cells were collected immediately and 6 mL of cells were incubated aerobically (~21% O_2_) for 15 min. Cells were collected as above. RNA isolation and on-column DNase treatment were completed using the RNeasy Mini Kit and RNase-Free DNase Kit, respectively (Qiagen). RNA was eluted in 50 μL RNase-Free water (Qiagen). A second DNase treatment was then performed on the samples (Invitrogen). Total RNA was quantified and purity was assessed using a Nanodrop 2000 Spectrophotometer (ThermoScientific). Libraries of cDNA were created using SensiFAST cDNA Synthesis Kit (Bioline). Control reactions without reverse transcriptase were conducted to confirm the absence of contaminating DNA in all samples. Quantitative real time polymerase chain reaction (qRT-PCR) was performed using SYBR Green Supermix (Applied Biosystems) on an Applied Biosystems 7500 Fast Real Time PCR System using the primers listed in **Error! Reference source not found.**. Relative gene expression was analyzed using the 2^−ΔΔCT^ method (Livak & Schmittgen, 2001) with *gapA* serving as the internal control (Rodriguez *et al.*, 2011).

### *In vivo* virulence assays

Virulence assays were performed using a rabbit model of infective endocarditis (Paik *et al.*, 2005). Specific pathogen-free New Zealand white rabbits weighing 2-4 kg were purchased from RSI Biotechnology and Charles River Laboratories. We allowed them to acclimate to the vivarium at least 7 days prior to inoculation. The rabbits were anesthetized and a 19-gauge catheter (BD Bioscience) was inserted through the right internal carotid artery past the aortic valve to cause minor damage. The catheter was trimmed, sutured in place, and remained in the artery for the entire experiment. The incision was closed with staples. *S. sanguinis* experimental strains were grown overnight in BHI at 1% or 6% O_2_, diluted 10-fold into fresh BHI, incubated for 3 h, sonicated, washed and resuspended in PBS. The inoculum was further diluted in PBS to obtain desired cell concentrations and 0.5 mL of combined culture was inoculated via intravenous injection into an ear vein on the second day after surgery. For studies only assessing the virulence of single manganese transport systems, each strain was inoculated at ~1 x 10^7^ CFU mL^−1^. For studies assessing combination mutants of multiple systems, WT was inoculated at ~5 x 10^6^ CFU mL^−1^ and each Δ*ssaACB* derivative strain was inoculated at ~1 x 10^8^ CFU mL^−1^. Spare inoculum culture was plated on BHI agar with appropriate antibiotics to confirm bacterial counts. At 20 h post-inoculation, rabbits were euthanized by intravenous injection of Euthasol (Virbac AH). Following removal of the heart, catheter placement was verified and vegetations were removed. Vegetations were homogenized with PBS, sonicated, diluted, and plated on BHI agar with appropriate antibiotics as above. The results were reported as recovered CFU per rabbit for each strain and normalized to inocula ratios. All animal procedures were approved by Virginia Commonwealth University Institutional Animal Care and Use Committee and complied with applicable federal and institutional guidelines.

### Protein modeling and depiction

Alignment of TmpA, BmtA, and ZIPB was performed with Clustal W in Geneious 11.1 (geneious.com). TMDs were based on the α-helices of the crystal structure of ZIPB (5TSA). Depiction of TmpA in 2D was generated in Protter (Omasits *et al.*, 2014) and modified to match the TMDs determined in the alignment. Protein models were built in SYBYL-X 2.0. The alignment of TmpA TMD III was adjusted to best fit ZIPB TMD III due to the differences in length, although a “bridge” still is present where a helix should be. The N-terminus and loop between TMDs III and IV were removed from the alignment for the model since they were not present in the crystal structure due to inherent disorder.

Water molecules and metal ions were removed from the ZIPB protein (PDB: 5TSA) and hydrogen atoms were retained. One hundred models were built, minimized with hydrogens over 10,000 iterations with a gradient of 0.5 and a Gastieger-Huckel charge. One model was chosen for further analysis with an RMSD of 1.783. Residue N173 was replaced with D and the model was again minimized as described above.

Positions within a cellular membrane were predicted using OPM (https://opm.phar.umich.edu/) and visualized in JMol 3.0 using FirstGlance (http://jmol.sourceforge.net/).

### Evolutionary analysis

Sequences of *S. sanguinis* SK36 TmpA and SsaB and *S. mutans* UA159 TmpA were used as queries for BLASTP searches of the Non-redundant protein or Refseq Select proteins database and the top hits were selected. For each species selected, only one sequence was used for the phylogenetic analysis. The evolutionary history was inferred using the Neighbor-Joining method (Saitou & Nei, 1987) with a gap opening penalty of 3.0 and a gap extension penalty of 1.8 for the multiple alignment stage (Hall, 2008). The evolutionary distances were computed using the Poisson correction method (Zuckerkandl & Pauling, 1965) and are in the units of number of amino acid substitutions per site. All ambiguous positions were removed for each sequence pair. Evolutionary analyses were conducted in MEGAX (Kumar *et al.*, 2018). 16S rRNA groups of streptococci, with the exception of *S. ratti*, were labeled based on Nobbs *et al.* (2009).

### Data analysis and presentation

Statistical tests were performed in GraphPad Prism (graphpad.com) and significance was determined by t-test or analysis of variance (ANOVA) as indicated in the figure legends. Tests were paired only if matching was effective. *P-*values ≤ 0.05 were considered significant. Graphs and figures were constructed using GraphPad Prism (graphpad.com) or Biorender (Biorender.com).

## Acknowledgments

We thank Ping Xu (VCU Philips Institute) for use of his strains and Jens Kreth and Nyssa Cullin (Oregon Health & Science University) for providing the IFDC *S. sanguinis* strain and corresponding protocol. We would like to thank Shannon Green, Seon-Sook An, Nicai Zollar, and Rachel Korba (VCU Philips Institute) for assistance with animal experiments. We are grateful for the advice and technical support of Joseph Turner (VCU Department of Chemistry) and the protein modeling assistance provided by Glenn Kellogg and Claudio Catalano (VCU Department of Medicinal Chemistry). We appreciate the membrane protein technical advice of David Eide (University of Wisconsin-Madison), Nick Noinaj (Purdue University), Jeannine Brady (University of Florida), Brian Kloss (Center on Membrane Protein Production and Analysis), and Sara Palmer. We would also like to sincerely thank Ross Belvin (VCU Philips Institute) for his advice and review of the manuscript.

## Author Contributions

TP and KK performed the experiments. TP and TK wrote the manuscript. All authors reviewed and approved the final version of the manuscript.

## Graphical Abstract

Depiction of manganese import into a *Streptococcus sanguinis* cell. The three known manganese-import systems are depicted and labeled as: ABC Transporter, ZIP-Family Protein, and NRAMP-Family Protein. Red arrows indicate the direction of manganese movement. Figure made with Biorender.

## Abbreviated Summary

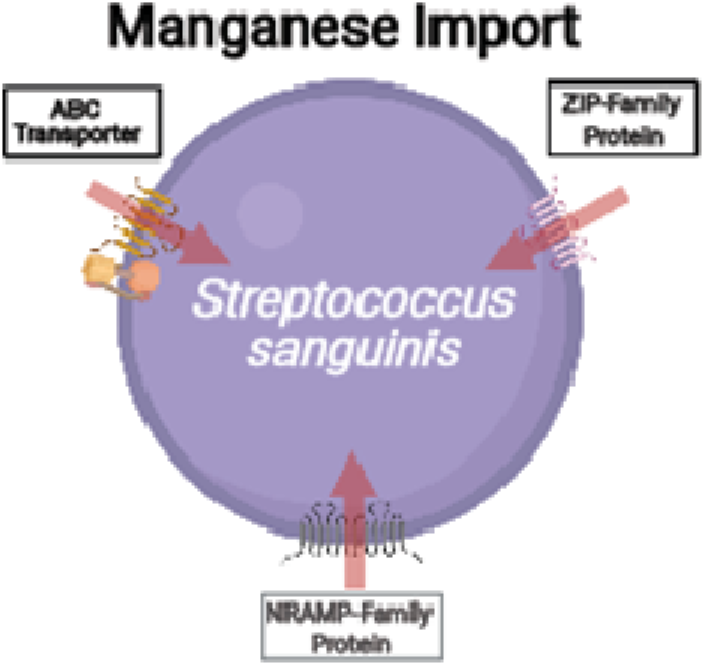

Manganese is an important metal for the growth in *Streptococcus sanguinis*, an oral bacterium that can cause a heart valve disease called infective endocarditis. Here we show that several different proteins transport manganese into *S. sanguinis* cells and have varying degrees of importance.

## Supporting Information

Puccio_Kunka_Kitten_Supp_Tables.xlsx Excel document Supplementary Tables

Puccio_Kunka_Kitten_Suppinfo.pdf PDF document Supplementary Figures

## Notes

**Funding Statement:** This work was supported by the National Institutes of Health: award F31 DE028468 to TP from the National Institute of Dental and Craniofacial Research and award R01 AI114926 to TK from the National Institute of Allergy and Infectious Diseases. The content is solely the responsibility of the authors and does not necessarily represent the official views of the National Institutes of Health.

**Conflict of Interest Disclosure:** Authors have no conflicts to disclose.

### Competing Interest Statement

The authors have declared no competing interest.

