## Supplementary Material for "Contribution of metal transporters of the ABC, ZIP, and NRAMP families to manganese uptake and infective endocarditis virulence in *Streptococcus sanguinis*"

### Supplemental Material

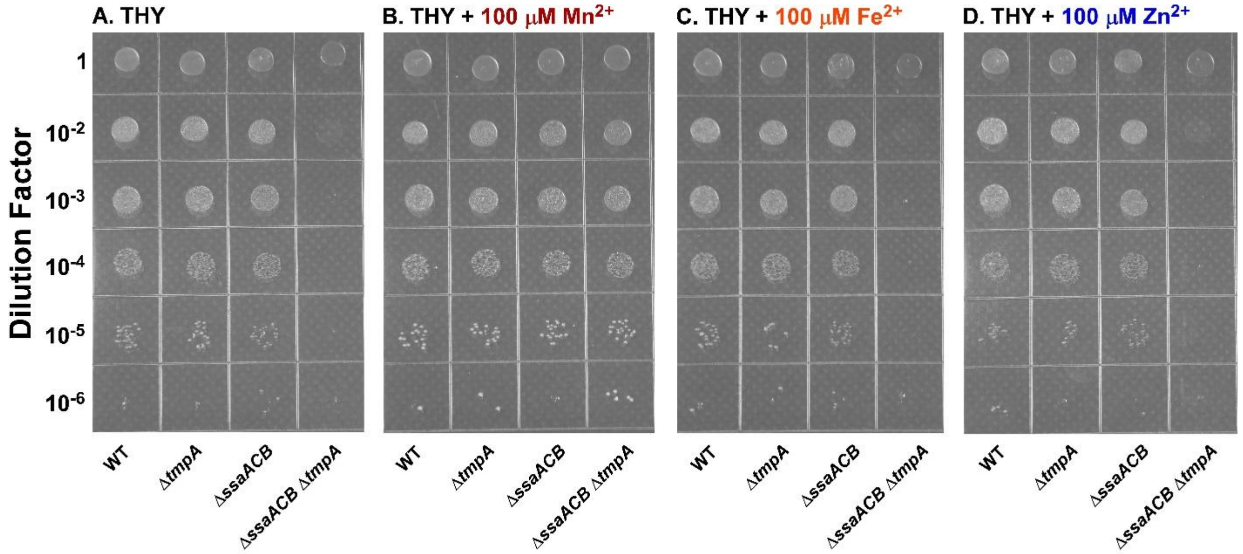


**Figure S1. Spot plating of WT, Δ*ssaACB*, and Δ*tmpA* strains**

Cultures were diluted as indicated and spotted in atmospheric conditions (~21% $O_2$) on plates of (A) THY, (B) + 100 µM $Mn^{2+}$, (C) + 100 µM $Fe^{2+}$, or (D) + 100 µM $Zn^{2+}$. The plates were incubated anaerobically for 24 h and imaged. Image representative of three replicates.

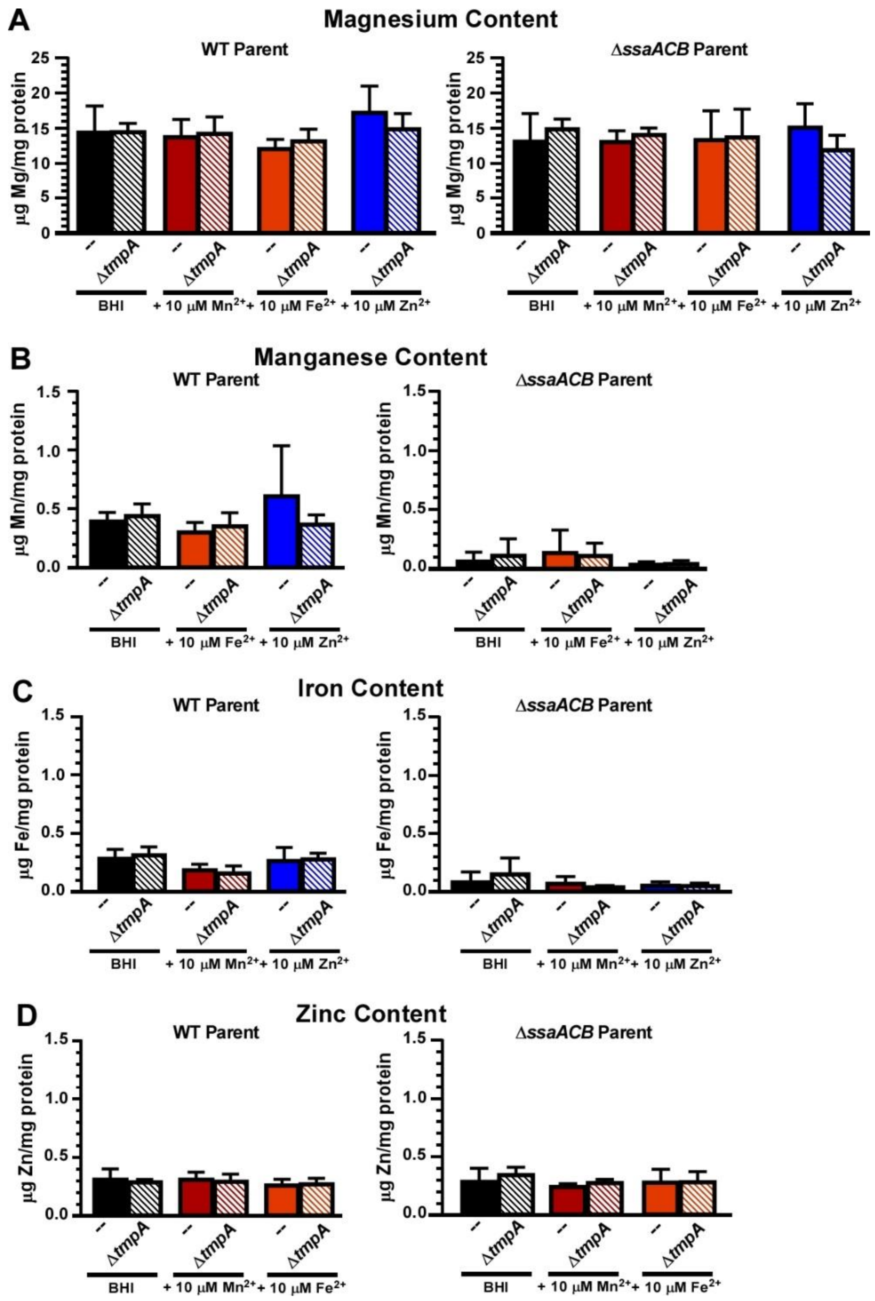


**Figure S2. Metal content of SK36 manganese-transporter mutants**

(A) Magnesium, (B) manganese, (C) iron, and (D) zinc content of each SK36 strain was assessed after growth in BHI ± 10 µM $Mn^{2+}$, $Fe^{2+}$, or $Zn^{2+}$ using ICP-OES and normalized to protein levels. Levels in BHI alone are the same as Figure 4 and are shown here for reference. Mutant strains generated from the WT parent are shown on the left and those made from the Δ*ssaACB* parent are on the right. Means and standard deviations of at least three replicates are displayed. Significance was determined by one-way ANOVA with Bonferonni's multiple comparisons test comparing each mutant and respective parent for each condition. No comparisons revealed significant differences.

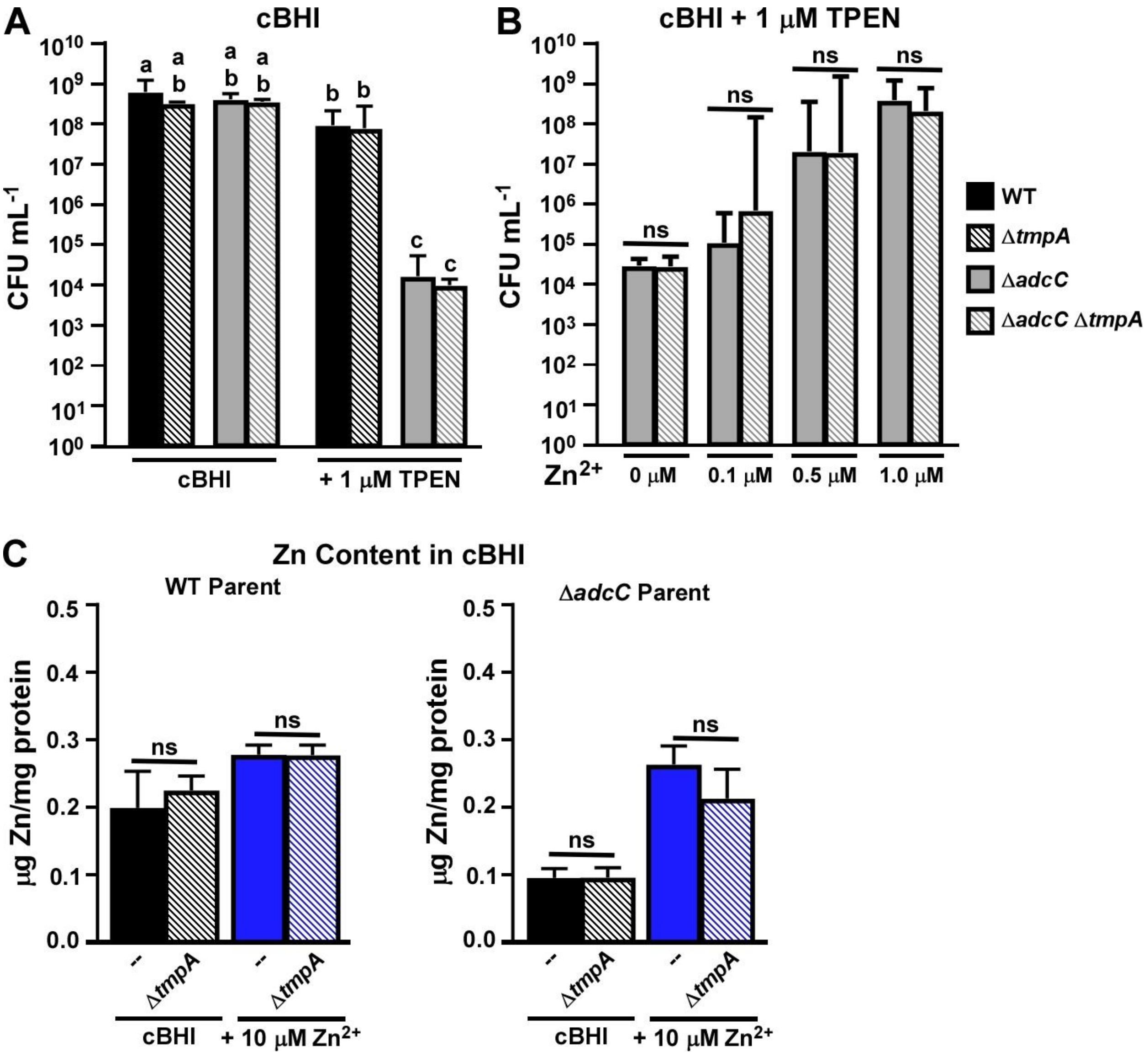


**Figure S3. Growth of metal transporter mutants in cBHI + TPEN at 1% $O_2$**

(**A**) Growth of the Δ*adcC* and Δ*tmpA* strains in cBHI ± 1 µM TPEN at 1% $O_2$ was measured. Means and standard deviations of at least 3 replicates are shown. Significance was determined by one-way ANOVA with a Tukey multiple comparisons post-test. Bars with the same letter are not significantly different from each other ($P > 0.5$) (**B**) Growth of each Δ*adcC* mutant in cBHI + 1 µM TPEN with added Zn was assessed. Means and standard deviations of at least 3 replicates are shown. Significance was determined by unpaired two-tailed t-tests comparing the Δ*adcC* Δ*tmpA* mutant to its Δ*adcC* parent under zinc concentration. (**C**) Cellular metal content of each strain was assessed after growth in cBHI using ICP-OES and normalized to protein levels. Means and standard deviations of three experiments are displayed. Significance was measured by one-way ANOVA with Bonferonni's multiple comparisons test comparing each mutant and respective parent for each condition.

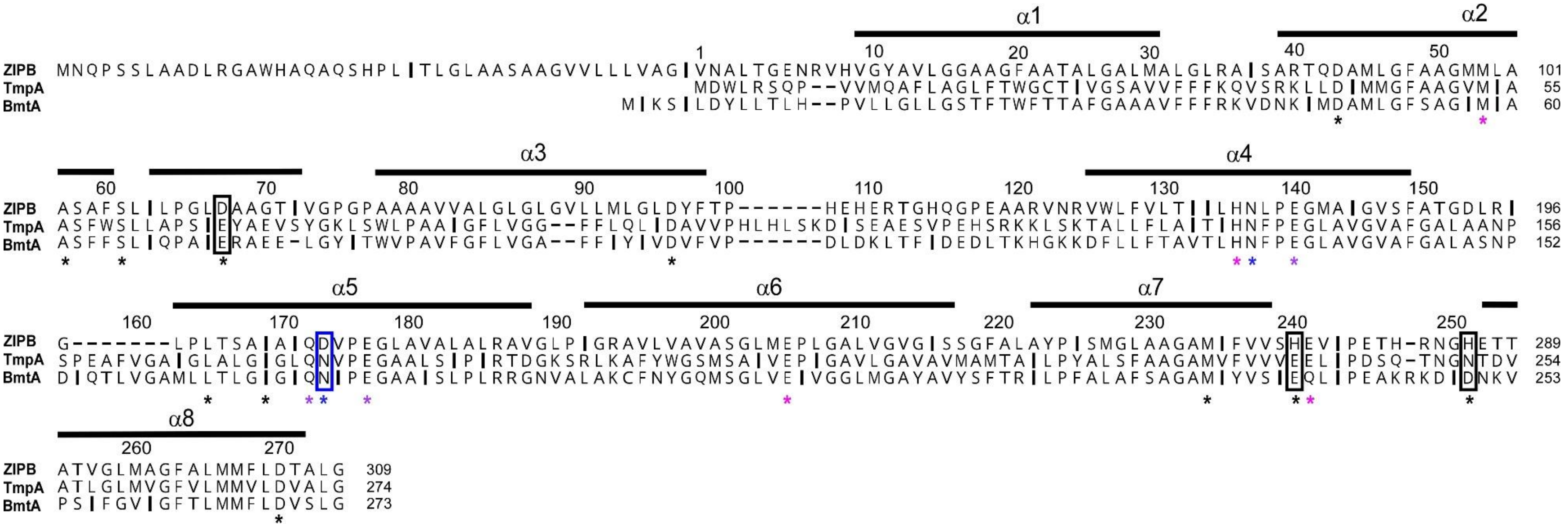


**Figure S4. Alignment of bacterial ZIP family proteins**

ZIP proteins from *B. bronchiseptica* (ZIPB), *S. sanguinis* (TmpA), and *B. burgdorferi* (BmtA) were aligned in Geneious. Numbers above sequences are based on TmpA; numbers on right indicate position for each strain. TMDs (α) are indicated by the horizontal lines and asterisks indicate metal binding residues in ZIPB (Zhang *et al.*, 2017). Magenta, blue, and purple asterisks indicate the residue binds to the metal at the M1 site, the M2 site, or both sites, respectively. Boxes indicate metal binding residues from ZIPB that are not conserved between ZIPB and the other two proteins.

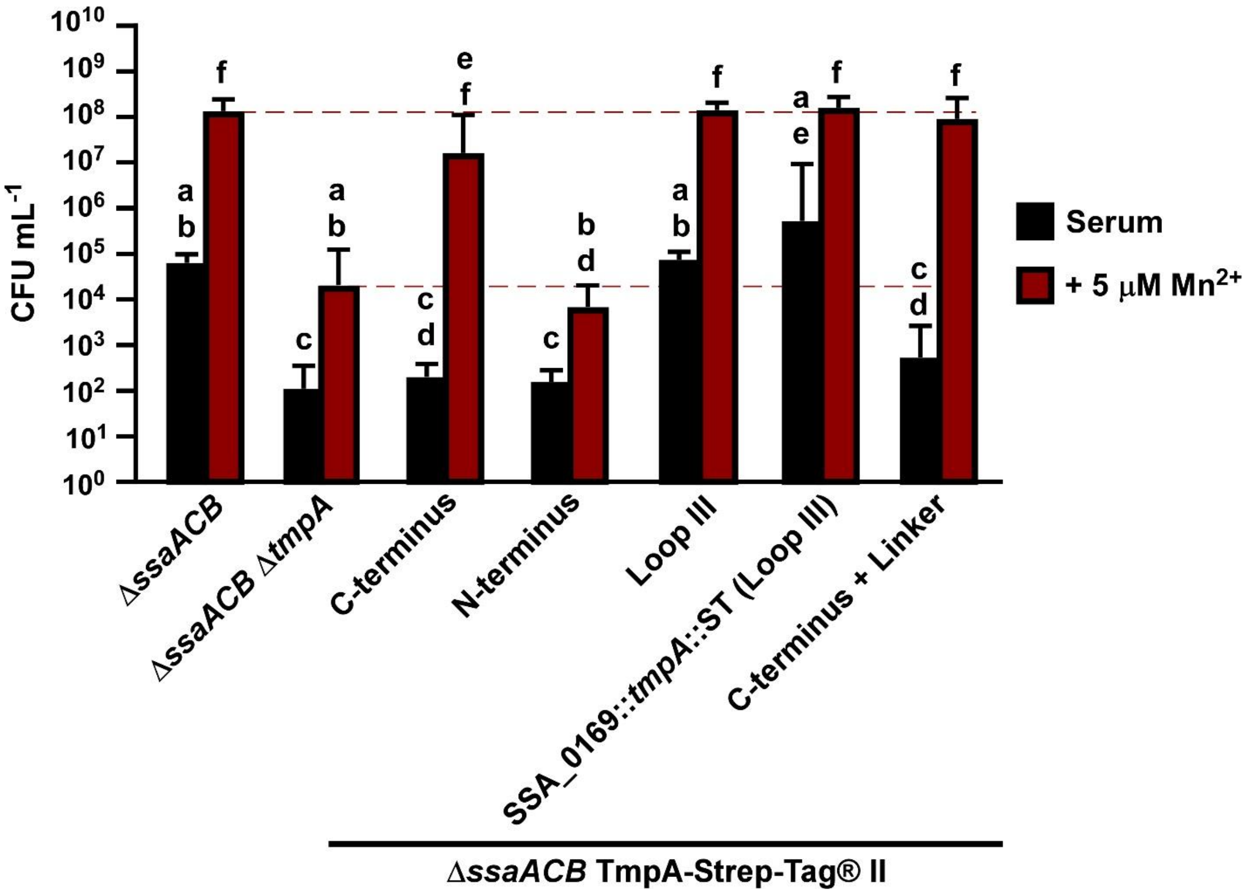


**Figure S5. Growth of Δ*ssaACB* TmpA-Strep-Tag® II mutants in serum**

Growth of various TmpA-Strep-Tag® II strains in the Δ*ssaACB* background were assessed in 6% $O_2$ serum ± 5 µM $Mn^{2+}$. Means and standard deviations of at least three independent experiments are shown. Significance was determined by one-way ANOVA with a Tukey multiple comparisons post-test. Bars with the same letter are not significantly different from each other ($P > 0.05$). Dashed lines indicate growth of the Δ*ssaACB* parent and Δ*ssaACB* Δ*tmpA* mutant in each condition for reference.

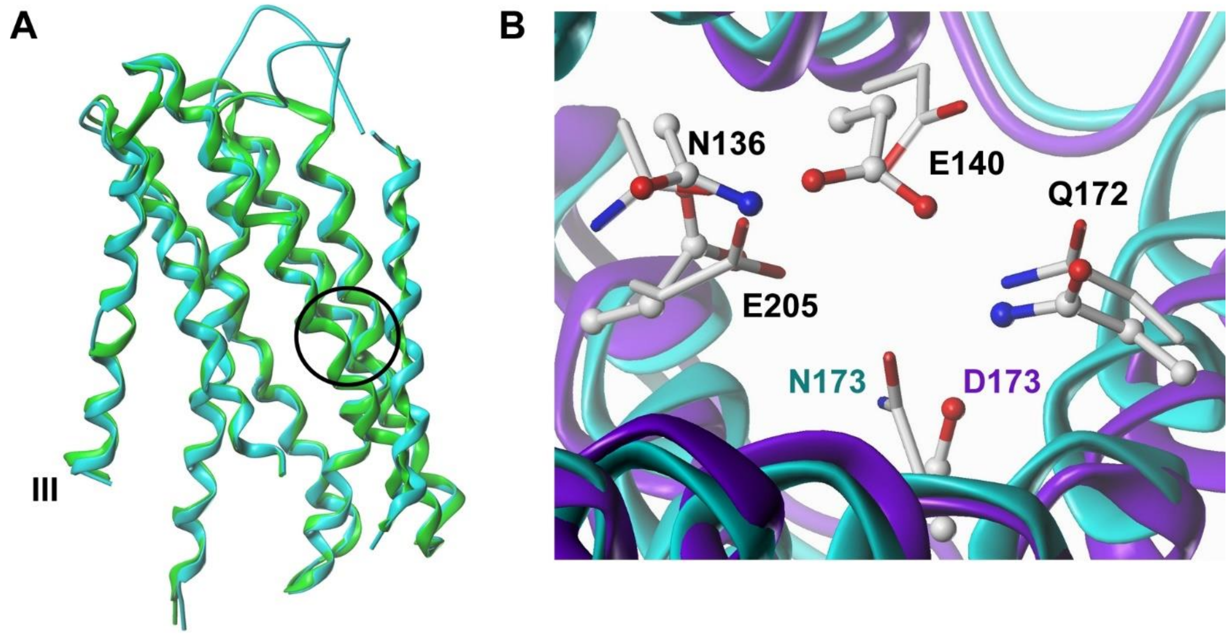


**Figure S6. Model of TmpA based on the crystal structure of ZIPB**
(A) TmpA model (cyan) based on the ZIPB crystal structure (PDB: 5TSA) (green). Loop III is indicated for reference. The circle indicates the region of the predicted "M2" metal binding site. (B) View of the predicted M2 site with WT TmpA (cyan; sticks) overlaid with the N173D mutant version (purple; ball and stick).

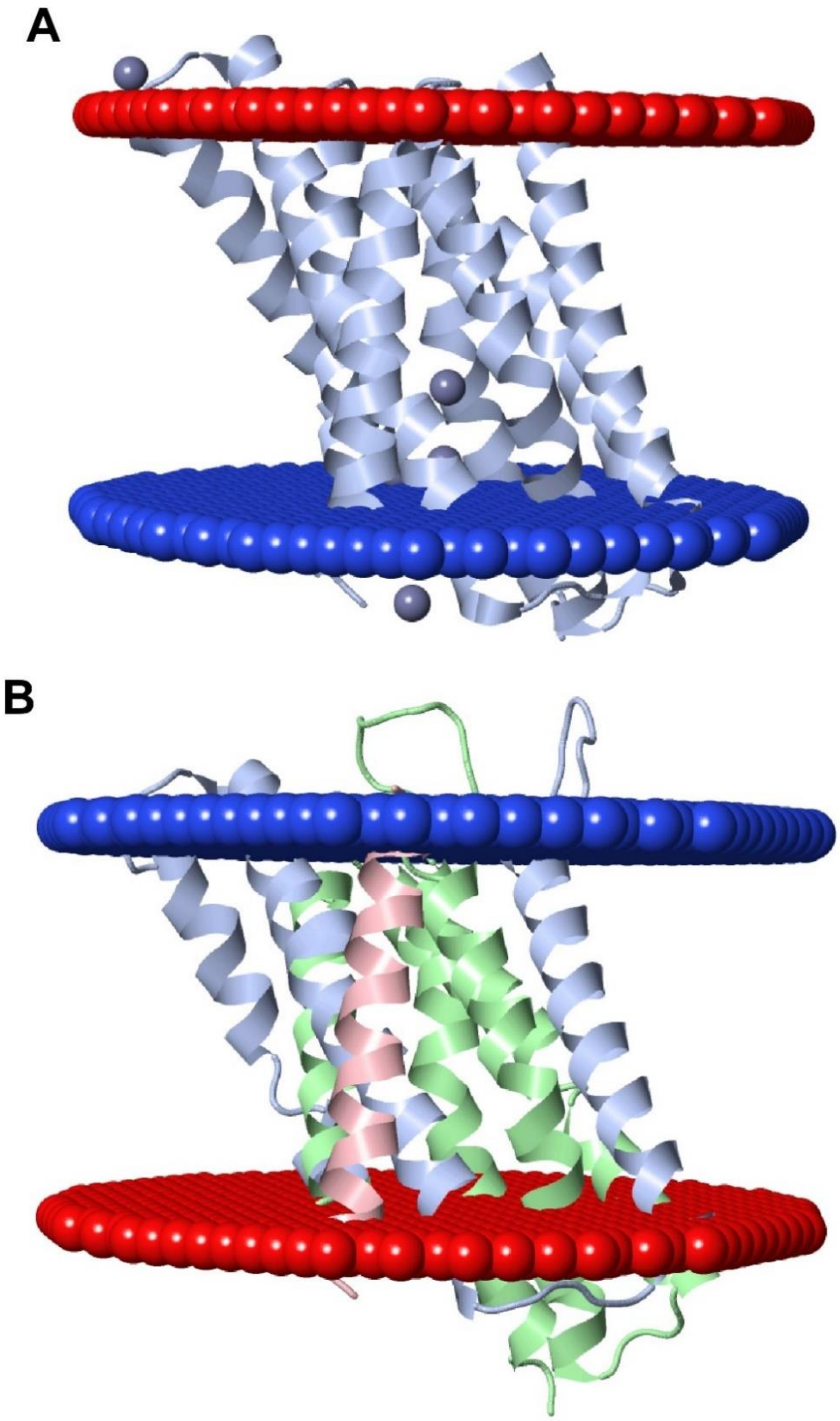


**Figure S7. Depiction of ZIP proteins within a membrane**
(A) The crystal structure of ZIPB (PDB: 5TSA) with zinc and cadmium ions and (B) the model of TmpA were predicted within the cellular membrane using OPM (https://opm.phar.umich.edu/) and visualized with FirstGlance in Jmol (http://jmol.sourceforge.net/). The top is the outward-facing side and the bottom is the cytoplasmic side.

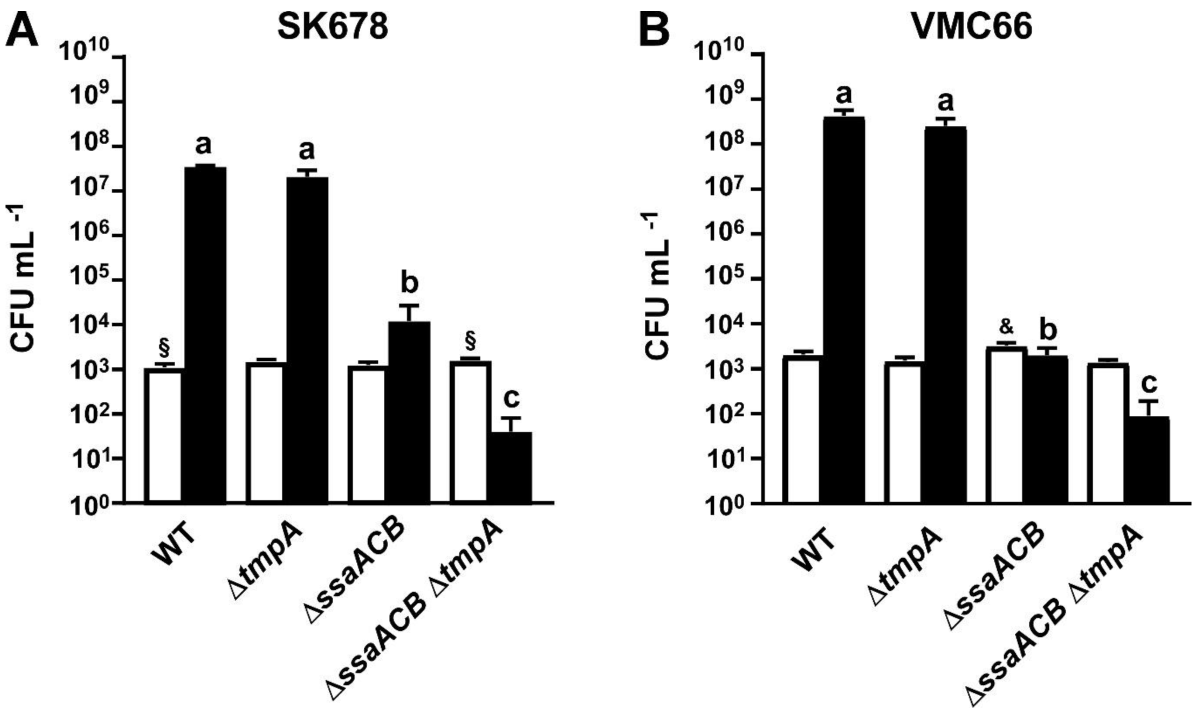


**Figure S8. Growth of Mn-transport mutants of SK678 and VMC66 in low-oxygen serum**
Growth of two *S. sanguinis* strains and their respective Mn-transporter mutants in serum at 1% $O_2$ for 24 h was assessed. Means and standard deviations of at least three independent experiments are displayed. Significance was assessed by one-way ANOVA with a Tukey multiple comparisons separately for $T_0$ and $T_{24}$ values. $T_{24}$ bars that share a letter within a chart are not significantly different from each other ($P > 0.05$). $T_0$ bars with **§** are significantly different from each other. The $T_0$ bar with **&** is significantly different from all other $T_0$ values in that chart.

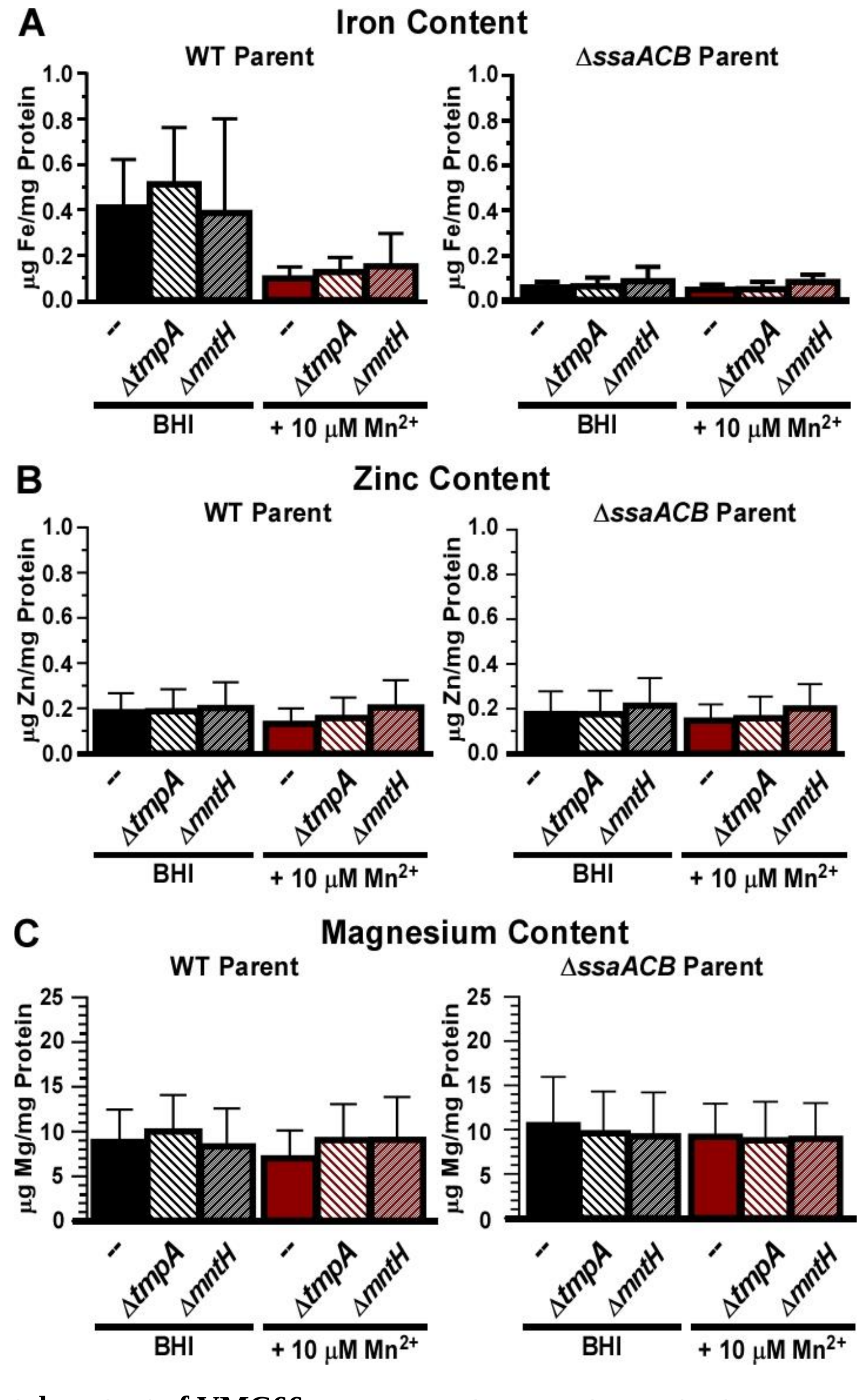


**Figure S9. Metal content of VMC66 manganese-transporter mutants**

(A) Iron, (B) zinc, and (C) magnesium content of each VMC66 strain was assessed after growth in BHI ± 10 µM $Mn^{2+}$ using ICP-OES and normalized to protein levels. Mutant strains generated from the WT parent are shown on the left and those made from the Δ*ssaACB* parent are on the right. Means and standard deviations of three experiments are displayed. Significance was determined by one-way ANOVA with Bonferonni's multiple comparisons test comparing each mutant and respective parent for each condition. No comparisons revealed significant differences.

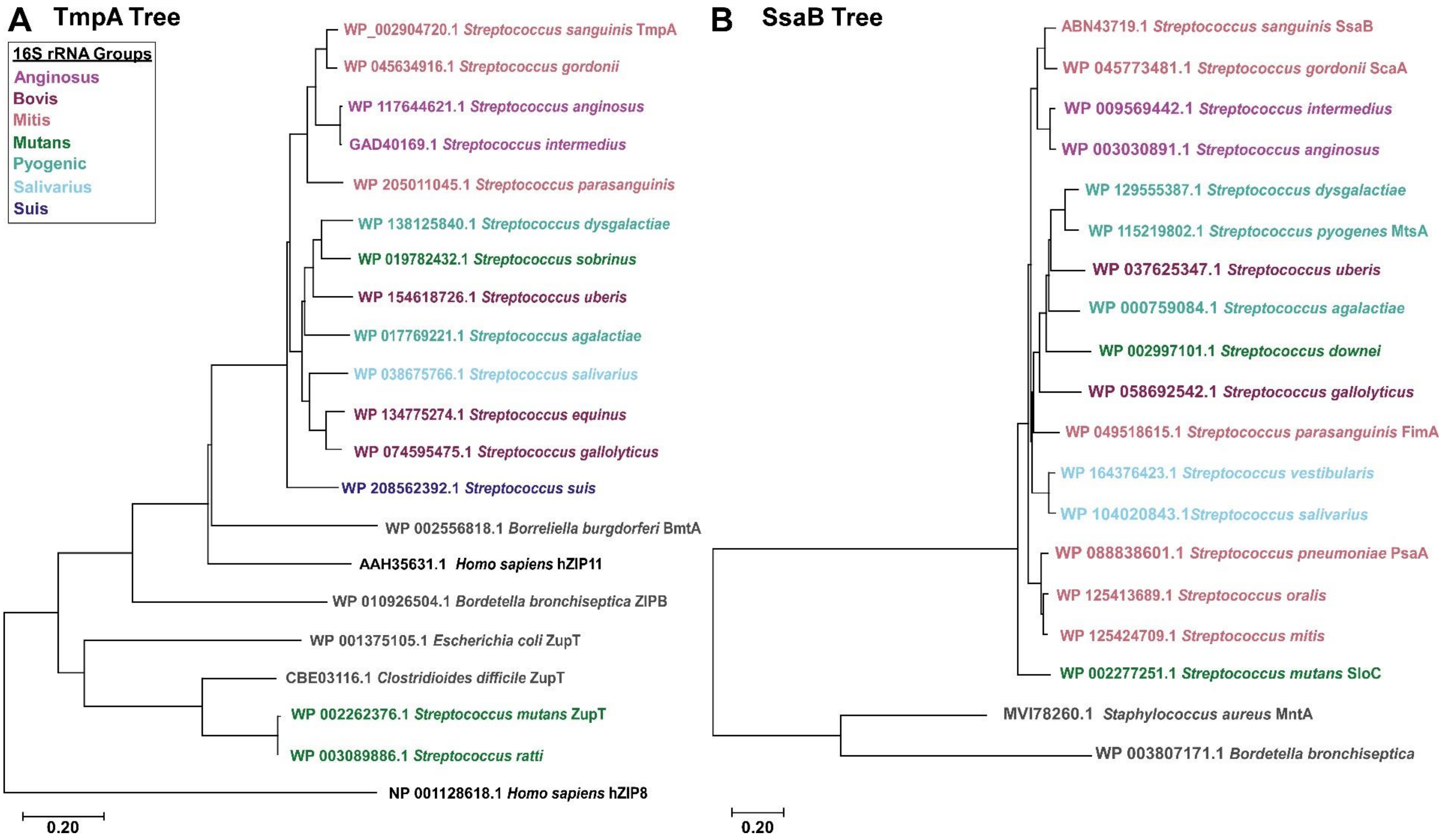


**Figure S10. Phylogenetic trees of *S. sanguinis* TmpA and SsaB homologs**

The evolutionary histories of selected top-scoring BLASTP matches to *S. sanguinis* SK36 (A) TmpA and (B) SsaB sequences were inferred. The optimal trees with the sum of branch length of (A) 5.04 and (B) 4.74 are shown. Streptococcal species are colored based on their 16S rRNA groups (Nobbs *et al.*, 2009); other bacterial species are listed in gray. Outgroups for each tree are (A) hZIP8 and hZIP11 (black) and (B) *B. bronchiseptica* and *S. aureus*. *C. difficile* ZupT from strain R20291 (Zackular *et al.*, 2020) was included in (A) for reference.

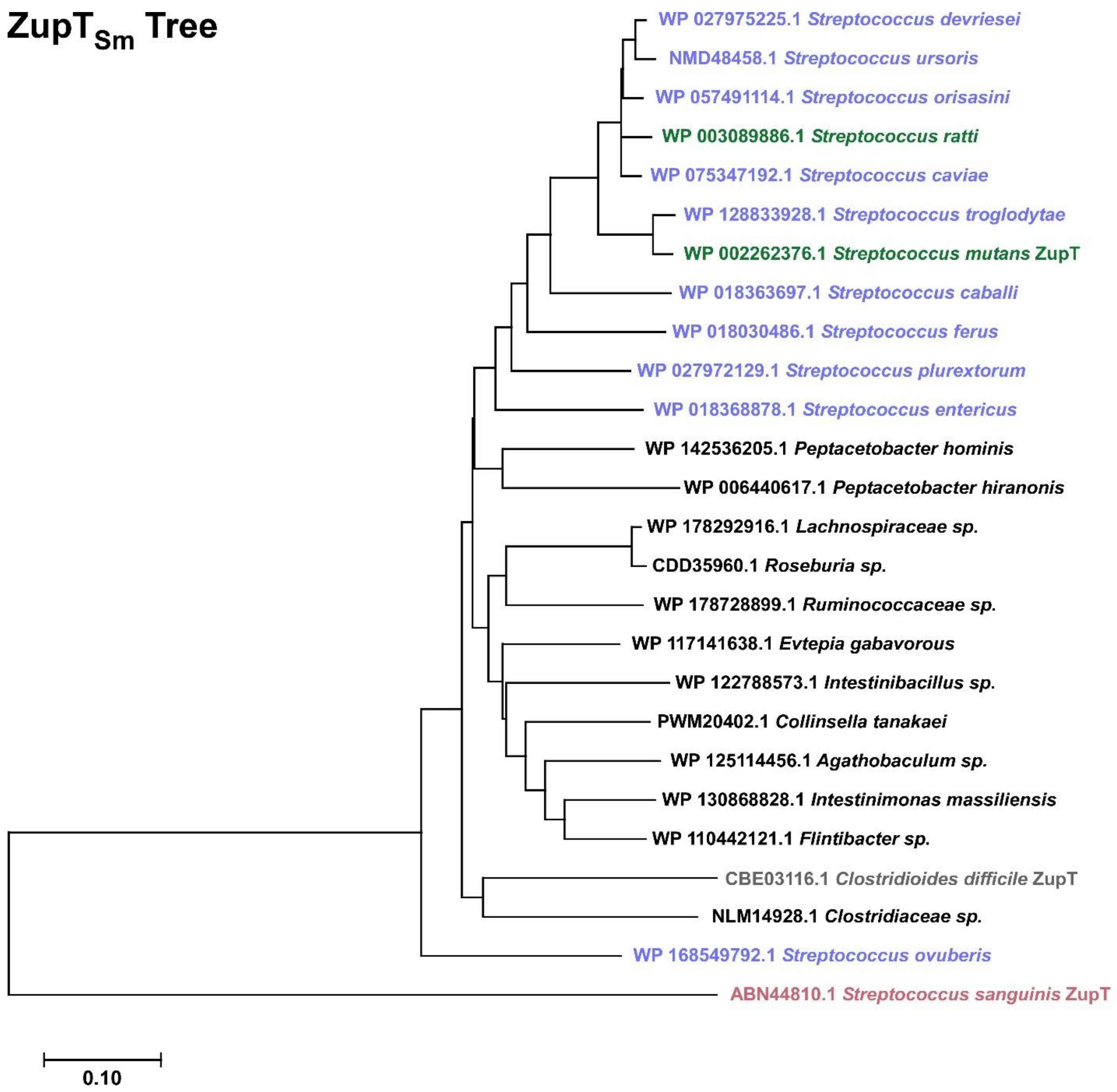


**Figure S11. Phylogenetic tree of *S. mutans* ZupT homologs**

The evolutionary history of the selected top-scoring BLASTP matches to the *S. mutans* UA159 ZupT sequence was inferred. The optimal tree with the sum of branch length of 3.35 is shown. Streptococcal species are colored; all other bacterial species are black. *S. sanguinis* TmpA (rose) was included as an outgroup. *C. difficile* ZupT (gray) was also included for reference.

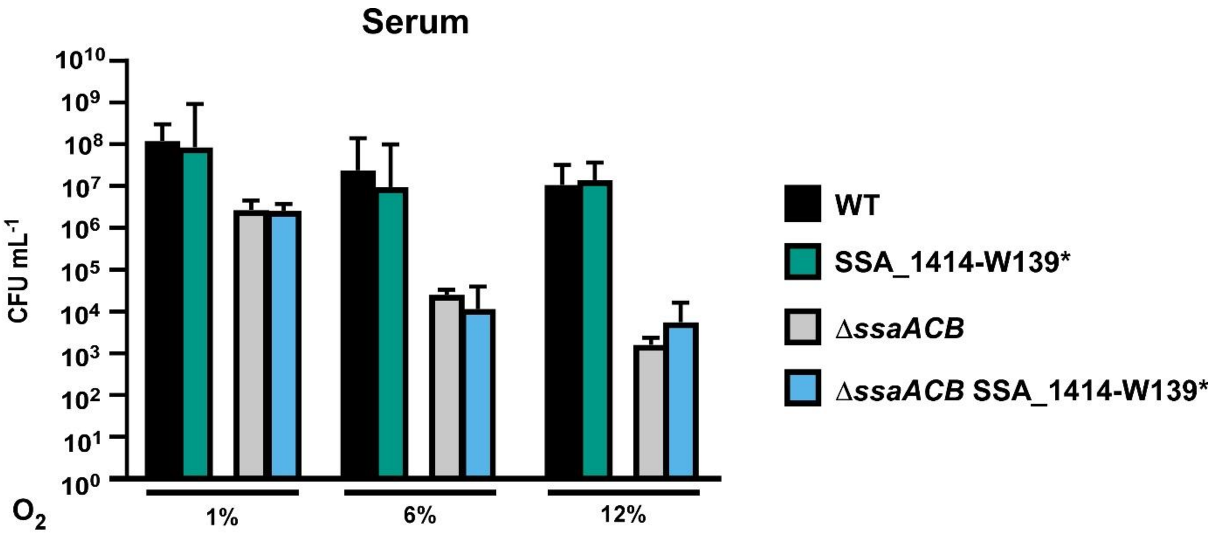


**Figure S12. Comparison of the growth of the SSA_1414-W139* mutants**
Cultures of each SSA_1414-W139* mutant as compared to their respective parent strain in serum at various $O_2$ concentrations. Means and standard deviations of three replicates are depicted. The strains were determined to be not significantly different from each other in each condition tested as determined by unpaired two-tailed t-test.

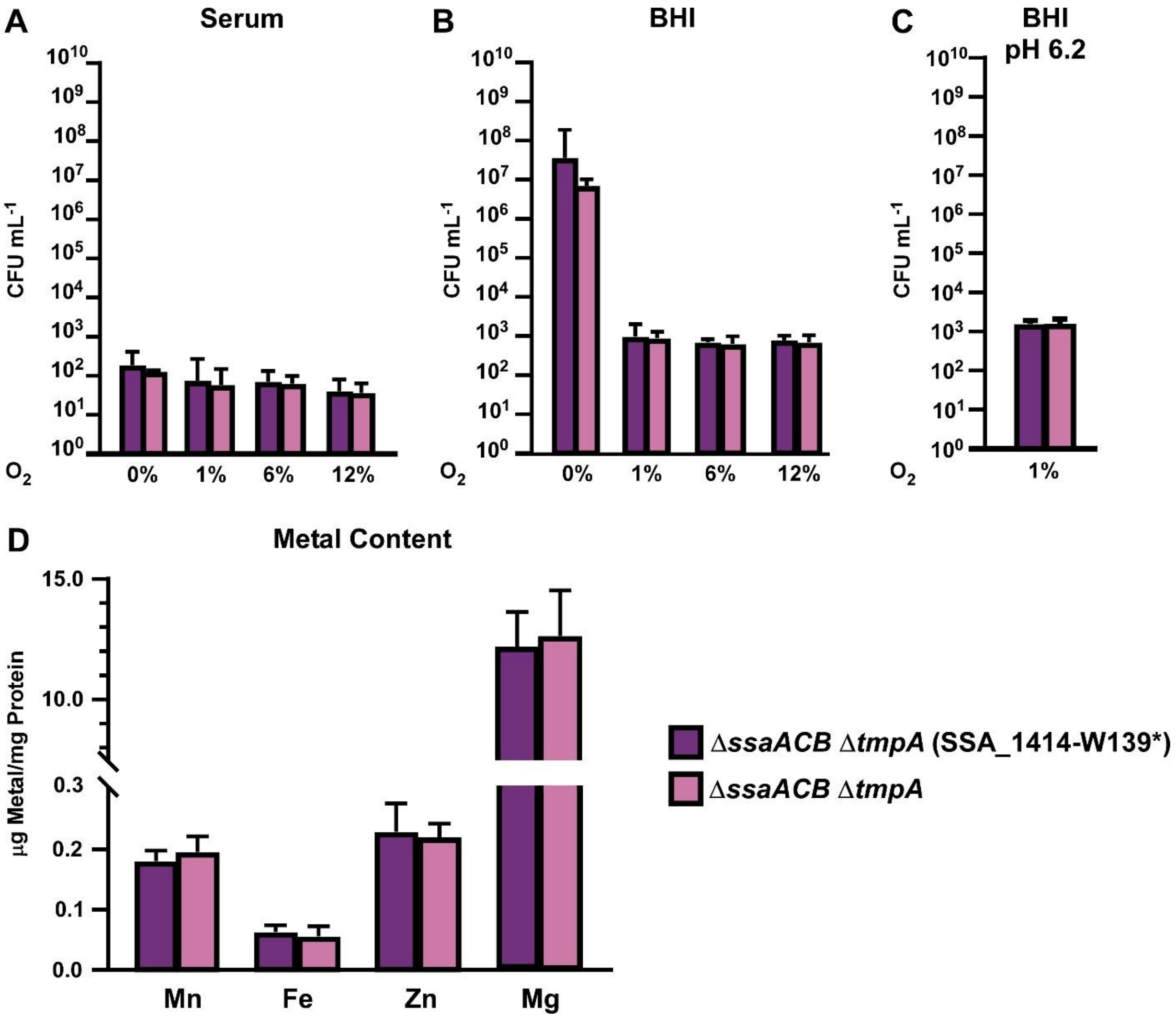


**Figure S13. Comparison of Δ*ssaACB* Δ*tmpA* mutants with and without SNP in SSA_1414.** Cultures of each Δ*ssaACB* Δ*tmpA* mutant in serum (**A**), normal BHI (**B**), or acidic BHI (**C**) were grown at the indicated $O_2$ concentrations. JFP227 is the original version and JFP377 is the new, clean version. Metal content of cells grown in atmospheric conditions (~21% $O_2$) in BHI with 10 µM $Mn^{2+}$ as measured by ICP-OES (**D**). Means and standard deviations of three replicates are depicted in each chart. The strains were determined to be not significantly different from each other in each condition tested as determined by paired two-tailed t-test.
